# Weakly supervised Unet: an image classifier which learns to explain itself

**DOI:** 10.1101/2022.09.09.507144

**Authors:** Robert John O’Shea, Carolyn Horst, Thubeena Manickavasagar, Daniel Hughes, James Cusack, Sophia Tsoka, Gary Cook, Vicky Goh

**Affiliations:** Department of Cancer Imaging, King’s College London, London, UK; Department of Radiology, Guy’s and St Thomas’ NHS Foundation Trust, London, UK; Department of Radiology, Liverpool University Hospitals NHS Foundation Trust, London, UK; Department of Natural and Mathematical Sciences, King’s College London, London, UK; King’s College London & Guy’s and St Thomas’ PET Centre, Guy’s and St Thomas’ NHS Foundation Trust, London, UK

**Keywords:** Explainable Artificial Intelligence, Model Interpretation, Weakly Supervised Learning, Lung Neoplasms, Tumour Segmentation;

## Abstract

**Background:** Explainability is a major limitation of current convolutional neural network (CNN) image classifiers. A CNN is required which supports its image-level prediction with a voxel-level segmentation.

**Methods:** A weakly-supervised Unet architecture (WSUnet) is proposed to model voxel classes, by training with image-level supervision. WSUnet computes the image-level class prediction from the maximal voxel class prediction. Thus, voxel-level predictions provide a causally verifiable saliency map for the image-level decision.

WSUnet is applied to explainable lung cancer detection in CT images. For comparison, current model explanation approaches are also applied to a standard CNN. Methods are compared using voxel-level discrimination metrics and a clinician preference survey.

**Results:** In test data from two external institutions, WSUnet localised the tumour precisely at voxel-level (Precision: 0.93 [0.93-0.94]), achieving superior voxel-level discrimination to the best comparator (AUPR: 0.55 [0.54-0.55] vs. 0.36 [0.35-0.36]). Clinicians preferred WSUnet predictions in most test instances (Clinician Preference Rate: 0.72 [0.68-0.77]).

**Conclusions:** WSUnet is a simple extension of the Unet, which facilitates voxel-level modelling from image-level labels. As WSUnet supports its image-level prediction with a causative voxel-level segmentation, it functions as a self-explaining image classifier.

Graphical Abstract
The weakly-supervised Unet converts voxel-level predictions to image-level predictions using a global max-pooling layer. Thus, loss is computed at image-level. Following training with image-level labels, voxel-level predictions are extracted from the voxel-level output layer.

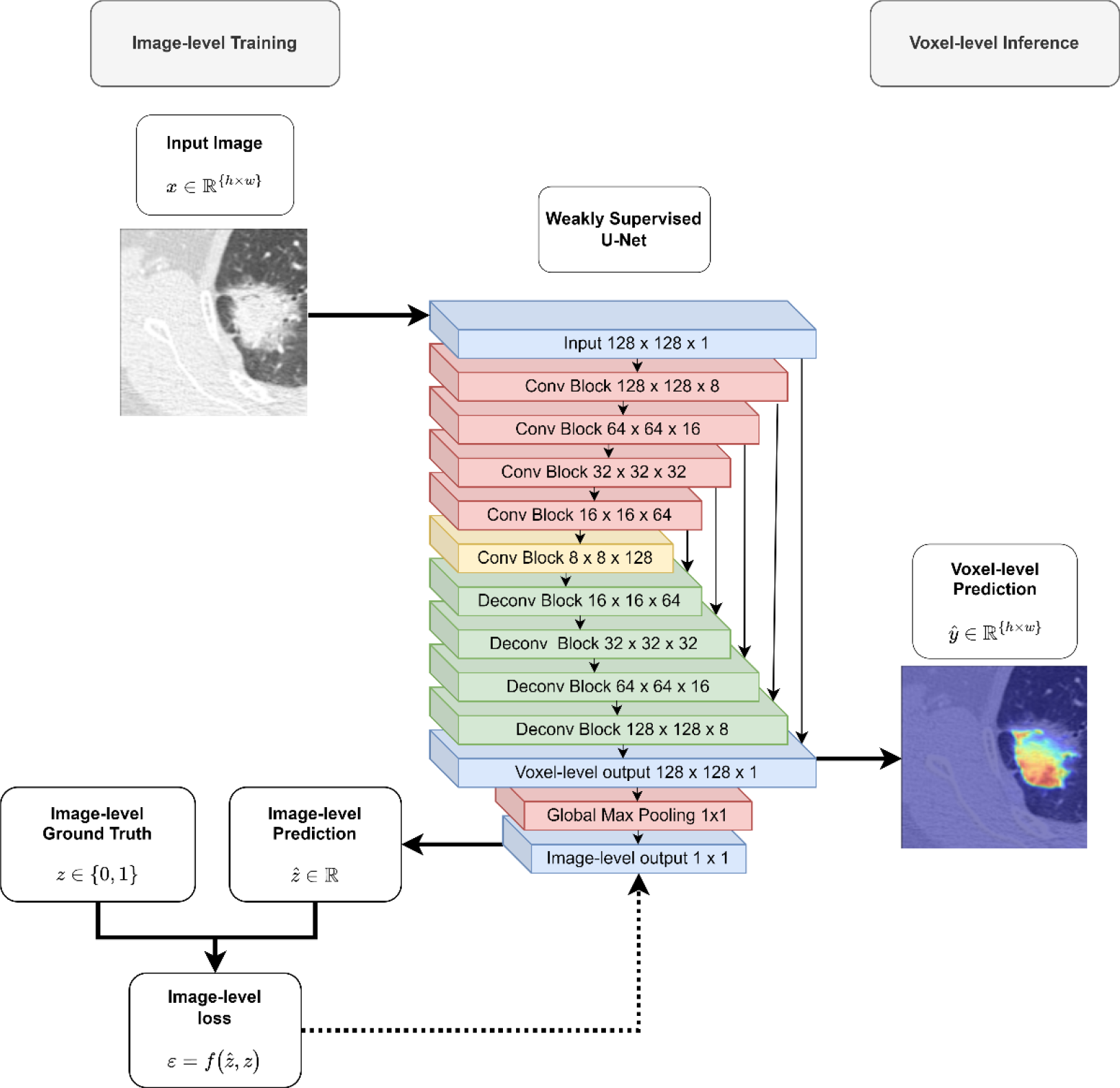

**Funding:** Authors acknowledge funding support from the UK Research & Innovation London Medical Imaging and Artificial Intelligence Centre; Wellcome/Engineering and Physical Sciences Research Council Centre for Medical Engineering at King’s College London [WT 203148/Z/16/Z]; National Institute for Health Research Biomedical Research Centre at Guy’s & St Thomas’ Hospitals and King’s College London; National Institute for Health Research Biomedical Research Centre at Guy’s & St Thomas’ Hospitals and King’s College London; Cancer Research UK National Cancer Imaging Translational Accelerator [C1519/A28682]. For the purpose of open access, authors have applied a CC BY public copyright licence to any Author Accepted Manuscript version arising from this submission.

**HIGHLIGHTS:**

1. WSUnet is a weakly supervised Unet architecture which can learn semantic segmentation from data labelled only at image-level.
2. WSUnet is a convolutional neural network image classifier which provides a causally verifiable voxel-level explanation to support its image-level prediction.
3. In application to explainable lung cancer detection, WSUnet’s voxel-level output localises tumours precisely, outperforming current model explanation methods.
4. WSUnet is a simple extension of the standard Unet architecture, requiring only the addition of a global max-pooling layer to the output.

## INTRODUCTION

Explainability is a well-known limitation of convolutional neural network (CNN) image classifiers, whose “black-box” nature presents various clinical issues (Amann et al., 2020; Grote and Berens, 2020; van der Velden et al., 2022). Traditionally, radiologists justify diagnoses with corresponding image findings, providing evidence which is independently verifiable. In contrast, CNN decisions are not explicitly justified. Consequently, it is difficult to verify that model has made an appropriate prediction, using relevant image features. Confounding, where images are classified based on spurious features, poses risks of misclassification, discrimination, and vulnerability to adversarial attacks in CNN models (Amann et al., 2020; Badgeley et al., 2019; Kaviani et al., 2022; van der Velden et al., 2022). For example, Badgeley demonstrated that a model predicting fracture in hip radiographs depended significantly upon patient characteristics and image acquisition parameters, failing to discriminate fractures from normal radiographs when these factors were controlled (Badgeley et al., 2019). The absence of justification for CNN decisions complicates interpretation by clinicians and patients alike, compromising responsibility, communication and capacity to consent (Amann et al., 2020; Grote and Berens, 2020). CNNs’ opacity also impedes their application to pathobiological discovery. In contrast, linear models’ transparency has elucidated mechanisms in fields such as genomics (Waldmann et al., 2013). A transparent CNN, which could reliably identify the region of interest motivating its decision, may identify new radiological disease characteristics. For these reasons, explainability is a central component of the European Commission’s Assessment List for Trustworthy Artificial Intelligence (European Commission, 2018), a key guideline for the prospective regulation of artificial intelligence development in high-risk applications such as healthcare.

As standard CNN decision functions are not easily invertible (Finzi et al., 2019), voxels’ contributions to the image-level predictions are unavailable, leading to the development of several indirect approaches to quantify surrogate voxel-importance measures (Ayyar et al., 2021), of which most may be classified as backpropagation-based and perturbation-based (van der Velden et al., 2022). Class activation mapping approaches (Selvaraju et al., 2020; Simonyan et al., 2014; Zhou et al., 2016) estimate regions’ importance by extracting gradients from the final convolutional layers. These methods are limited by low resolution and uncertain causality. The deconvolutional neural network (Zeiler and Fergus, 2014) aims to reverse engineer CNN decisions by passing each layers’ filter outputs to an inverted CNN pyramid, where upsampling and convolutional filtering map the image-level output to the voxel-level domain. Estimation of voxel-level importance is complicated by uninvertible operations such as max-pooling. Furthermore, as image-level predictions are independent of voxel-level outputs, explanations are not causally guaranteed. Voxel trainable attention (Jetley et al., 2018; Oktay et al., 2018) layers apply sigmoid activations to internal convolutional layers, so that the model may learn to deactivate irrelevant image regions. Although trainable attention provides an explanation which is causally relevant, interpolation is required to map the outputs to the voxel-level domain.

Perturbation-based methods measure voxels’ contributions through comparison of model predictions on modified and unmodified images. Occlusion sensitivity (Zeiler and Fergus, 2014) estimates regions’ importance by systematic occlusion and re-inference. Gradient integration (Sundararajan et al., 2017) estimates voxels’ importance through repeated inference on a series of progressively whitened images, allowing estimation of voxel-level gradients. Perturbation-based methods’ interpretability may be limited by the CNN decision functions’ nonlinearity and nonmonotonicity.

Conceptually, the problem of explainable image classification is closely related to that of weakly-supervised semantic segmentation, where voxel-level classification labels, *y* ∈ {0,1}^*h*×*w*^ are modelled from data labelled at image-level, *z* ∈ {0,1}. Fully convolutional neural networks (FCNs), have enjoyed success in fully supervised semantic segmentation tasks (Ronneberger et al., 2015; Siddique et al., 2021), where voxel-level labels are available. However, voxel annotation is laborious, requiring specialist expertise in the radiological setting (Mongan et al., 2020). Furthermore, human subjectivity or error complicate the use of manually-annotated labels, introducing variability into downstream outputs, such as radiomic features (Bianconi et al., 2021; Haarburger et al., 2020). Weakly-supervised approaches aim to train with incomplete labels, such as bounding boxes, point clouds, and image-level class labels. Of these, image-level labels are preferable (M. Zhang et al., 2020), as they are easily annotated and provide the weakest level of supervision.

Several weakly supervised semantic segmentation methods have been proposed, many using complex architectures and training schemas (M. Zhang et al., 2020). Zhang introduced a multi-stage system, in which a segmentation model was trained to predict pseudo masks derived from class activation mapping saliency maps and seed expansion (D. Zhang et al., 2020). Chaudry developed a fully convolutional attention network which integrated class activation mapping and convolutional attention mapping, erasing salient regions hierarchically (Chaudhry et al., 2017). Wei employed erasing to extend attention maps, iteratively retraining the model (Wei et al., 2017). Redondo-Cabrera proposed a dual network: a “hide-and-seek” network to obscure image regions and attempt image classification; and a “segmenter” network to apply the optimal “hiding” patterns learn object segmentations (Redondo-Cabrera et al., 2019). Li et al proposed a guided attention mapping network, extracting voxel-level predictions from internal attention layers and subsequently training these layers in a fully supervised manner (Li et al., 2018). Arun proposed a one-step classifier composed of a noisy unet in which the conditional distribution of each voxel could be estimated with respect to the class (Arun et al., 2020). Wang and Chen both addressed the issue of causality in class activation maps by integrating the class activation map into the class prediction, thereby restructuring the causal chain (Chen et al., 2022; Wang, 2022).

The objective of this proof-of-concept study was to develop a simple image classifier which provides a causally verifiable voxel-level explanation. A weakly-supervised FCN architecture is proposed, composed of a Unet with a global max-pooling layer appended to the output (WSUnet). WSUnet architecture is demonstrated in **Figure 1**. It was hypothesised that WSUnet could learn voxel-level classification from data labelled at image-level. The framework was applied to detect and localise non-small cell lung cancer (NSCLC) in computed tomography (CT) images from multiple centres.

**Figure 1.**
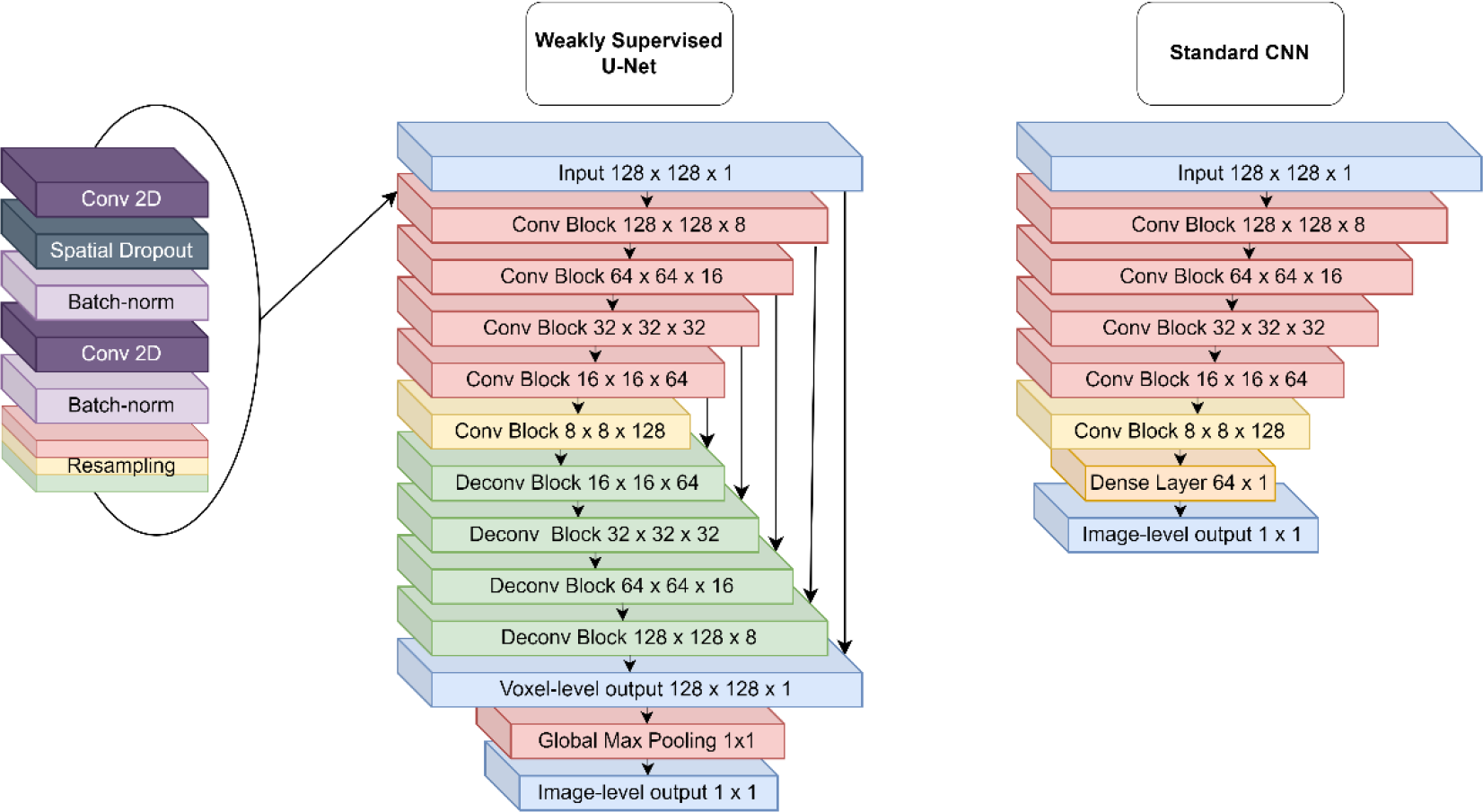
Weakly Supervised U-net and Standard CNN architectures. Blue layers represent inputs and outputs. Red, green and yellow layers represent downsampling, upsampling, and non-resampling convolutional blocks, respectively. Purple, charcoal and lilac layers represent convolutional, dropout and normalisation layers, respectively. Orange layers represent dense layers.

## MATERIALS AND METHODS

This study was performed in accordance with the CLAIM guidelines (Mongan et al., 2020). The CLAIM checklist is provided in **Supplementary Table 1.**

### Experimental Data

WSUnet was evaluated by application to NSCLC detection and localisation, using data from The Cancer Imaging Archive (Clark et al., 2013). Model development was performed with the Aerts et al. dataset (Aerts et al., 2019, 2014), which contains annotated retrospective CT data from 421 inoperable, pathologically confirmed, stage I-IIIb NSCLC patients at Maastricht University Medical Centre. Testing was performed in the Bakr et al. dataset (Bakr et al., 2017; Gevaert et al., 2012), which contains annotated retrospective CT data from 142 early stage NSCLC patients referred for surgical management at Stanford School of Medicine and the Veterans Association Hospital Palo Alto. Further information on the study data is provided in **Supplementary Information**. Subjects were excluded if CT images with tumour segmentations were unavailable, or if the gross tumour volume was not clearly identifiable in the annotation region labels.

### Preprocessing

CT images were converted to Hounsfield Units and scaled by a factor of 0.001. Patches of dimension 128 × 128 × 1 were sampled from axial CT slices. In each image, 16 patches were sampled from the tumour region and 16 were sampled from random centrepoints.

### Model Development

A weakly supervised Unet (WSUnet) was constructed by appending a global max-pooling layer to the output of a Unet with five convolutional and four deconvolutional blocks (487k trainable parameters). For comparison, a standard CNN model (SCNN) was generated with equivalent architecture (1,344k trainable parameters). Model architectures are described in **Figure 1.** Model weights were randomly initialised. Rectified linear activation was applied to hidden layers and sigmoid activation to the output layers. Spatial dropout was applied with a rate of 0.25. The models were optimised using the Adam optimiser with a learning rate of 0.001 using binary cross-entropy loss at the image level. Training and validation partitions were created from a random patient-disjoint 80-20 split of the Aerts dataset. Training and validation images were randomly augmented with horizontal flips, vertical flips and rotations by 90, 180 and 270 degrees. A single training run continued until validation loss plateaued for 5 epochs. Model training was performed with Tensorflow version 2.4.0, Keras version 2.2.4 and keras-unet (Chollet, François and et al., 2021; Google Inc., 2021; Zak, 2020). All code required to reproduce the results of this analysis is provided at https://github.com/robertoshea/wsss/.

### Model Testing

Test performance was evaluated in the Bakr dataset. Voxel-level outputs were extracted from the WSUnet model. **Figure 2** provides a schema of model training and testing. Four model interpretation methods were applied to the SCNN model using modified code from the tf-explain library (Meudec, 2021). GradCAM heatmaps were extracted from the seventh convolutional layer (GradCAM (16,16,64)) and ninth convolutional layer (GradCAM (8,8,128)). Nearest-neighbour interpolation was applied to scale the grad-cam outputs to the input dimension (128 × 128). Occlusion sensitivity was applied with an occlusion width of 10 voxels. Integrated gradients were measured with 10 whitening steps. Voxel-level NSCLC discrimination was evaluated using area under the precision-recall curve (AUPR). As WSUnet returns class probability estimates, calibration was also computed in terms of precision, recall and dice score, discretising by a threshold of 0.5. As the other methods do not generate class probabilities, calibration metrics were not computable. Image-level classification was also evaluated with accuracy, sensitivity, specificity and area-under the receiver operating characteristic curve (AUC) metrics.

**Figure 2.**
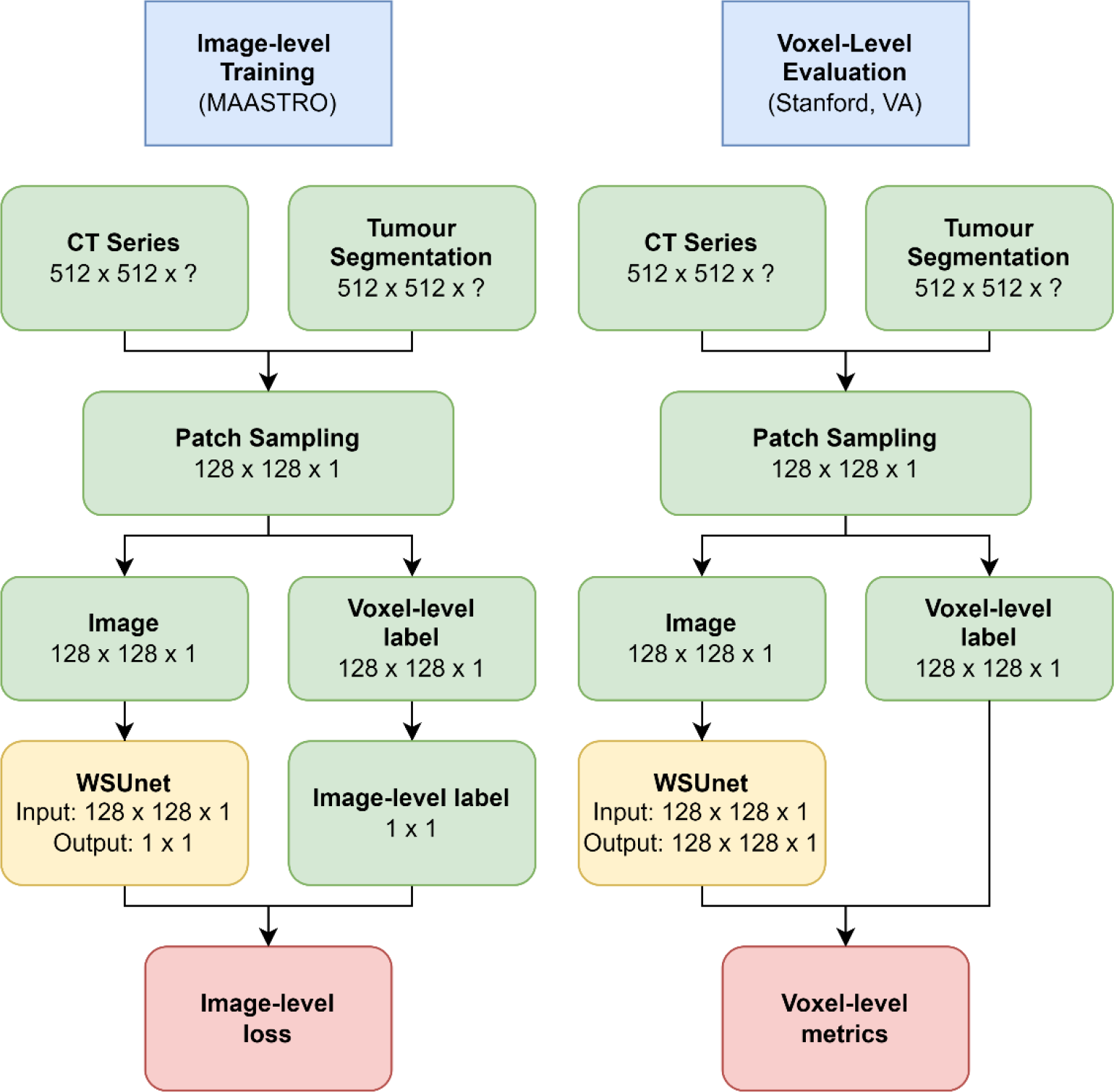
Model Training and Evaluation Schema.

Subjective clinical evaluation of voxel-level outputs was also performed by clinicians with subspecialty experience in thoracic cancer imaging, including one staff radiologist (VG), two specialist radiology residents (CH, JC) and two specialist oncology residents (TM, DH). One hundred images with tumour present were extracted from the test set, and each voxel-level prediction method applied. Clinicians were provided with input images, ground truth and a blinded, random ordering of each methods’ voxel-level predictions.

Clinicians selected the method which they considered to “best represent the tumour” in each image. Clinician preference rate was calculated as the frequency with which clinicians preferred the method, excluding instances in which they considered all methods uninformative. Clinicians also rated the tumour detection difficulty in each image (1: “tumour obvious”, 2: “tumour difficult to identify”, 3: “Tumour not visible in this image”). The clinician preference survey is provided in **Supplementary Data 1**.

Metric distributions were estimated with 500 nonparametric bootstraps and 95% confidence intervals for all metrics were estimated according to the 2.5^th^ and 97.5^th^ centiles of the bootstrap distribution.

## THEORY

Let *X* ∈ ℝ^*n*×*h*×*w*^ be an array of *n* greyscale images with height *h* and width *w*. Let *T* be a target object which may appear in the images. Let 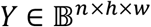 represent the corresponding array of binary voxel-level segmentation labels, such that that *Y*_*i*,*j*,*k*_ indicates the presence of target object in the *i*th image in the *j*th row, *k*th column voxel, *X*_*i*,*j*,*k*_.

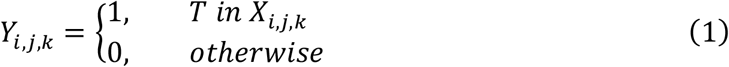

Let 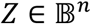 indicate the corresponding image-level labels, such that:

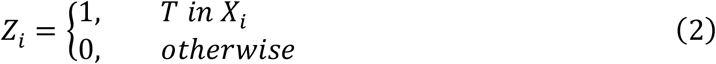

As a positive image is characterised by the presence of at least one positive voxel, the image-level label may be defined by the maximum voxel-level label, such that:

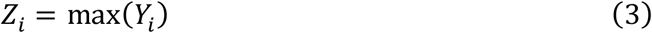

Therefore, nonzero image-level error implies nonzero voxel-level error, such that:

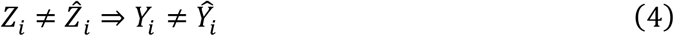

However, image-level correctness does not necessarily imply voxel-level correctness:

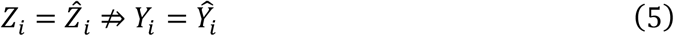

In the typical fully supervised setting, a voxel-level classification function 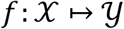 is learned using a dataset 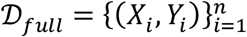. Model error 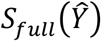 is quantified directly from a loss function 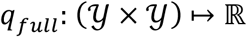 such that:

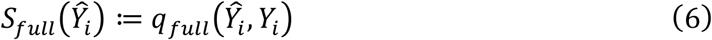

In the proposed weakly-supervised setting, *f* must be learned using a dataset of image-level labels 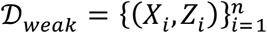. The max operator may be applied to voxel-level predictions to convert them to image-level, allowing application of an image-level loss function 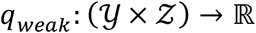 to create a weak objective 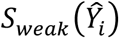 such that:

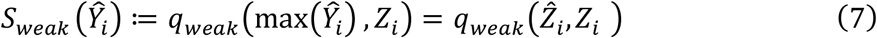

Thus, *q*_*weak*_ is fully supervised at image-level and weakly supervised at voxel-level. Hence, the proposed model learns a compound function 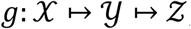, such that:

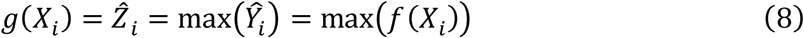

Therefore 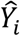 is a causal ancestor of 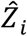. Following image-level training of *g*, the final global max operation may be removed, yielding the voxel-level prediction function, *f*. Thus, both 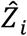 and 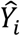 may be recovered. As 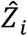 is a monotonic increasing function of 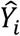, it provides easily interpretable voxel-level justification for the image-level decision.

## RESULTS

### Study data characteristics

Clinical characteristics of the training and testing cohorts are described in **Table 1**. Image acquisition parameters are provided in **Supplementary Table 1**. We excluded one subject from the Aerts dataset as the gross tumour volume (GTV) label could not be identified definitively, and 69 subjects were excluded from the Bakr dataset due to absence of segmentation data.

**Table 1.**
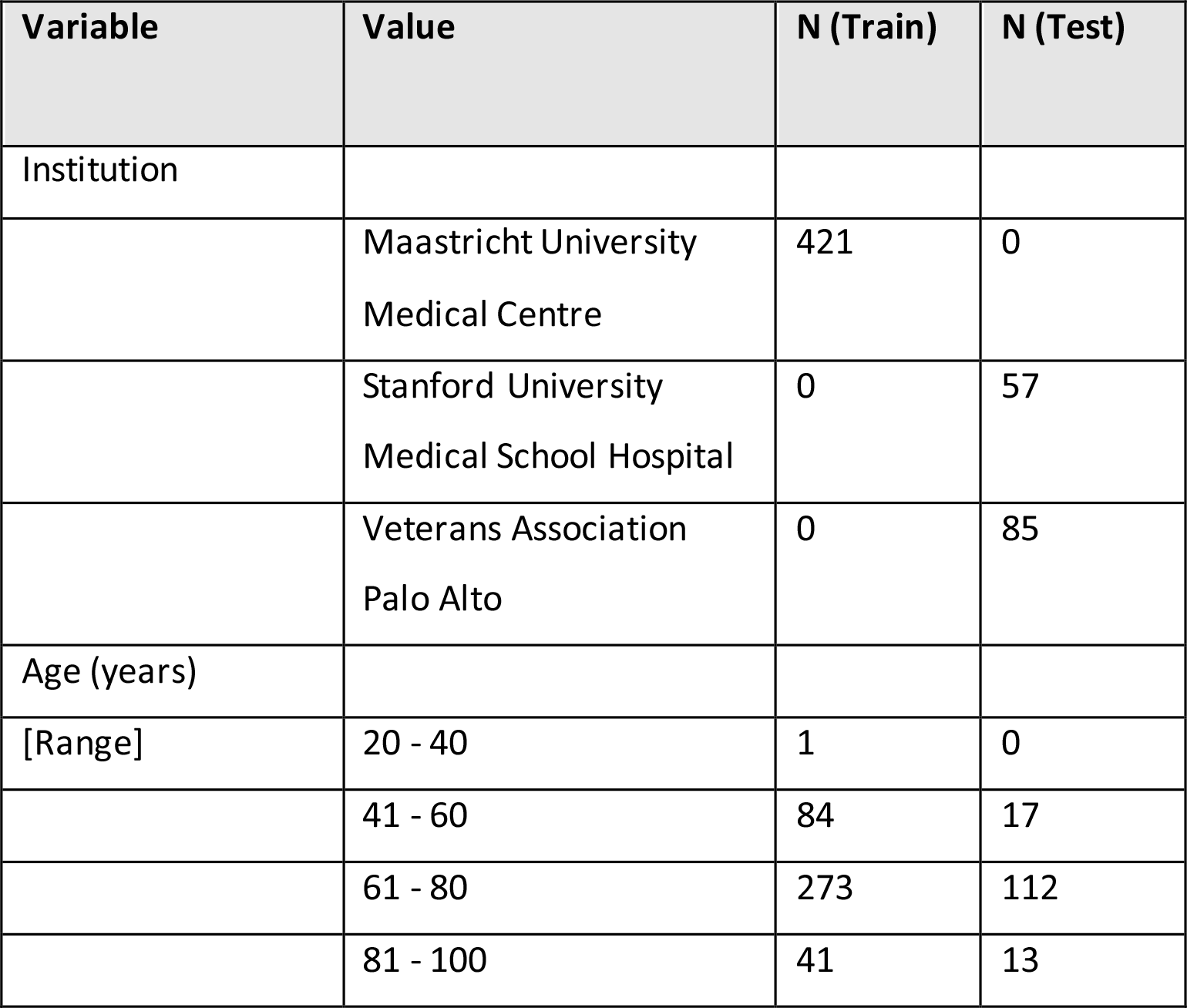
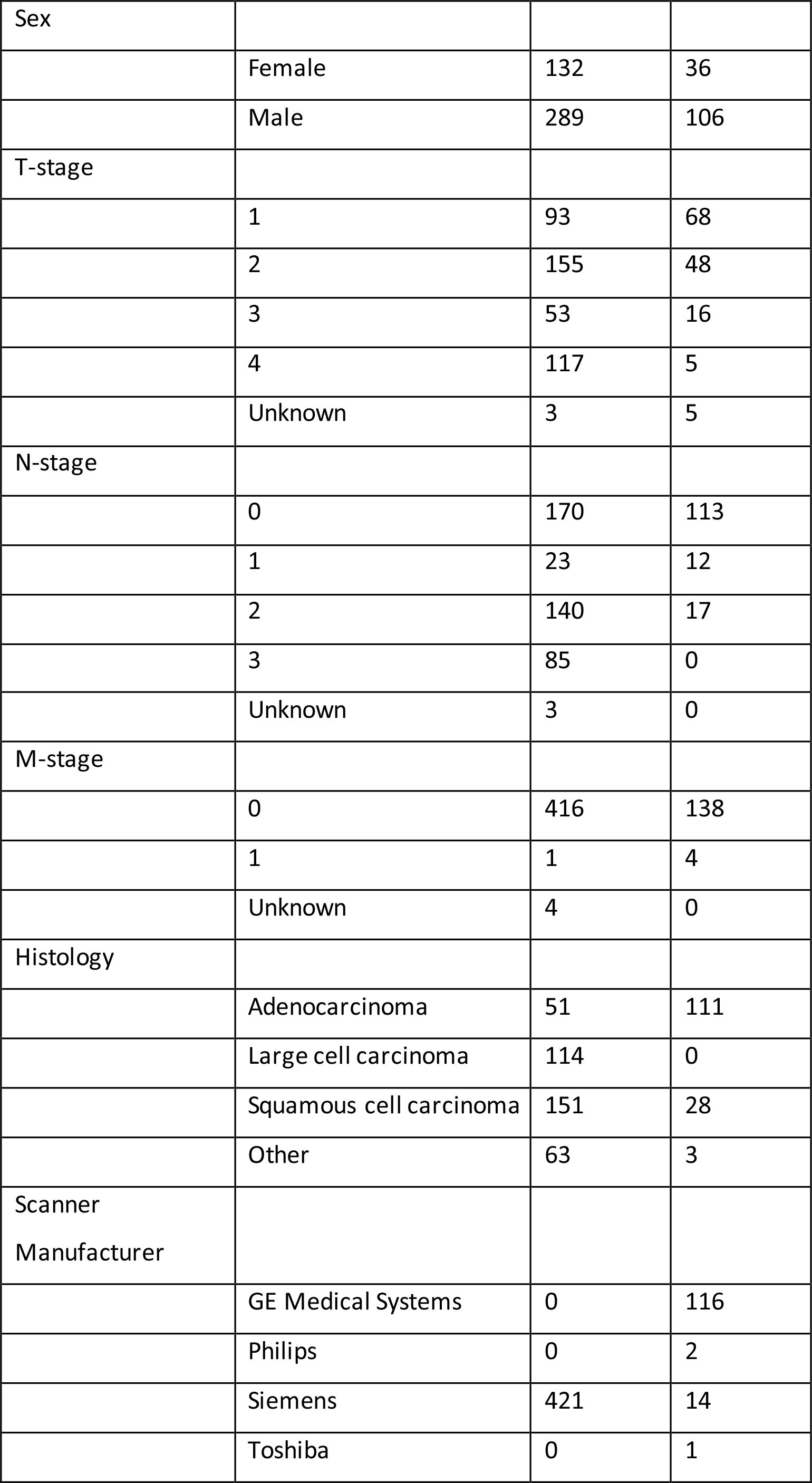
Clinical characteristics of the study population. Stage represents clinical stage in the training data and pathological stage in test data.

| Variable | Value | N (Train) | N (Test) |
| --- | --- | --- | --- |
| Institution |  |  |  |
|  | Maastricht University<br>Medical Centre | 421 | 0 |
|  | Stanford University<br>Medical School Hospital | 0 | 57 |
|  | Veterans Association<br>Palo Alto | 0 | 85 |
| Age (years) |  |  |  |
| [Range] | 20 - 40 | 1 | 0 |
|  | 41 - 60 | 84 | 17 |
|  | 61 - 80 | 273 | 112 |
|  | 81 - 100 | 41 | 13 |
| Sex |  |  |  |
|  | Female | 132 | 36 |
|  | Male | 289 | 106 |
| T-stage |  |  |  |
|  | 1 | 93 | 68 |
|  | 2 | 155 | 48 |
|  | 3 | 53 | 16 |
|  | 4 | 117 | 5 |
|  | Unknown | 3 | 5 |
| N-stage |  |  |  |
|  | 0 | 170 | 113 |
|  | 1 | 23 | 12 |
|  | 2 | 140 | 17 |
|  | 3 | 85 | 0 |
|  | Unknown | 3 | 0 |
| M-stage |  |  |  |
|  | 0 | 416 | 138 |
|  | 1 | 1 | 4 |
|  | Unknown | 4 | 0 |
| Histology |  |  |  |
|  | Adenocarcinoma | 51 | 111 |
|  | Large cell carcinoma | 114 | 0 |
|  | Squamous cell carcinoma | 151 | 28 |
|  | Other | 63 | 3 |
| Scanner<br>Manufacturer |  |  |  |
|  | GE Medical Systems | 0 | 116 |
|  | Philips | 0 | 2 |
|  | Siemens | 421 | 14 |
|  | Toshiba | 0 | 1 |

Clinicians considered positive instances moderately difficult to identify, assigning difficulty levels of “Tumour difficult to identify” and “Tumour not visible in this image” to 25.8% and 6% of test images, respectively. In many cases, ground glass changes were the only visible finding. Clinician preferences are provided in **Supplementary data 2**.

### Objective Performance Metrics

Voxel-level classification (segmentation) performance in test data is provided in **Table 2**. WSUnet’s voxel-level outputs localised NSCLC regions precisely (Precision: 0.93 [96% CI: 0.93-0.94]). Voxel-level recall was low (Recall: 0.16 [95%CI: 0.16-0.17]), indicating that the model classified images as positive without attending to the whole tumour region. WSUnet demonstrated strong discrimination at voxel-level (AUPR: 0.55 [95%CI: 0.54-0.55]), significantly outperforming the closest alternative, GradCAM (8,8,128) (AUPR: 0.36 [95%CI: 0.35-0.36]).

**Table 2.**
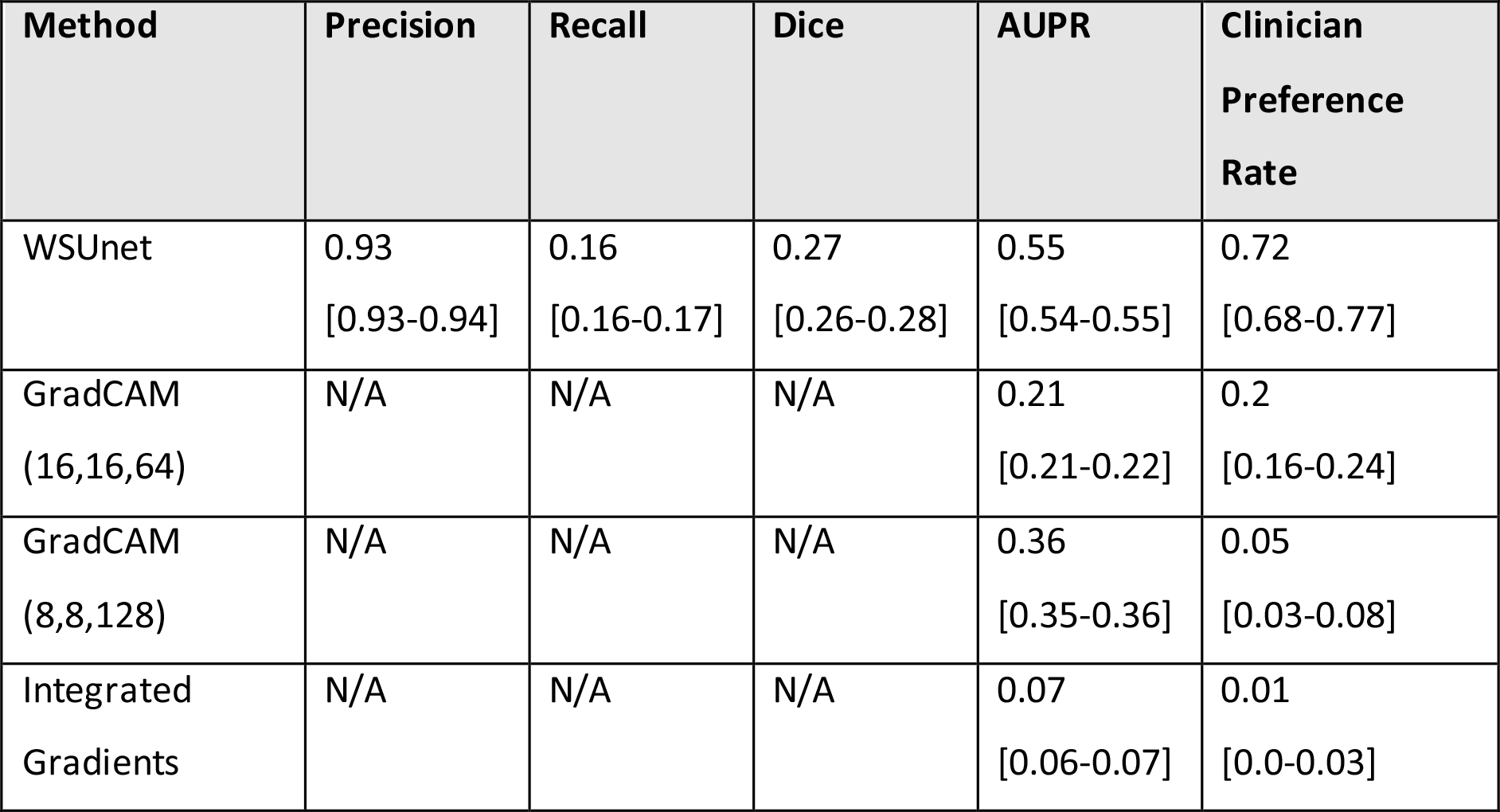
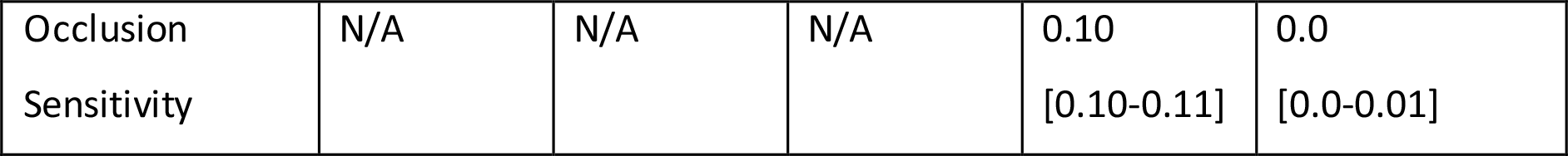
Voxel-level NSCLC classification results on test instances. Mean values and 95% confidence intervals are provided. Calibration metrics were not computable for GradCAM, Integrated Gradients and Occlusion Sensitivity methods. Clinician preference rate was calculated as the frequency with which clinicians preferred the method in 100 test instances, excluding instances in which they considered no model informative.

Image level classification results are provided in **Table 3**. WSUnet demonstrated similar image-level classification performance (Accuracy: 0.82 [0.81-0.82]; AUC: 0.91 [0.90-0.91]) to SCNN (Accuracy: 0.84 [0.84-0.85]; AUC: 0.91 [0.90-0.92]).

**Table 3.** Image-level NSCLC classification results on test instances. Mean values and 95% confidence intervals are provided.

| Method | Accuracy | Sensitivity | Specificity | AUC |
| --- | --- | --- | --- | --- |
| WSUnet | 0.82<br>[0.81-0.82] | 0.71<br>[0.7-0.73] | 0.95<br>[0.95-0.95] | 0.91<br>[0.9-0.91] |
| SCNN | 0.84<br>[0.84-0.85] | 0.79<br>[0.77-0.8] | 0.91<br>[0.91-0.92] | 0.91<br>[0.91-0.92] |

### Clinicians’ Performance Assessment

Clinicians strongly preferred WSUnet’s voxel-level outputs to current explainability methods. Excluding instances where no method was considered informative (26%), WSUnet outputs were preferred in 72% of test instances. GradCAM (16,16,128) and GradCAM (8,8,64) outputs were preferred in 20% and 5% of test instances, respectively. Integrated gradients and occlusion sensitivity outputs were preferred in fewer than 1% of test instances.

Methods’ voxel-level outputs are provided in **Figure 3**. Inspection of WSUnet’s voxel-level output confirms use of tumoural and peritumoural voxels to generate positive image-level classifications. Although “hot” regions were highly specific to tumour-related areas, several small nodules were missed. GradCAM outputs offered higher sensitivity to small tumours, however, rGradCAM (16, 16, 64) marked several ribs as “warm” and the resolution of GradCAM (8,8,128) outputs was low. Integrated gradient outputs surrounded the tumour region reliably – however positive regions were neither continuous nor specific. Occlusion sensitivity outputs were uninformative, differing minimally between inputs.

**Figure 3.**
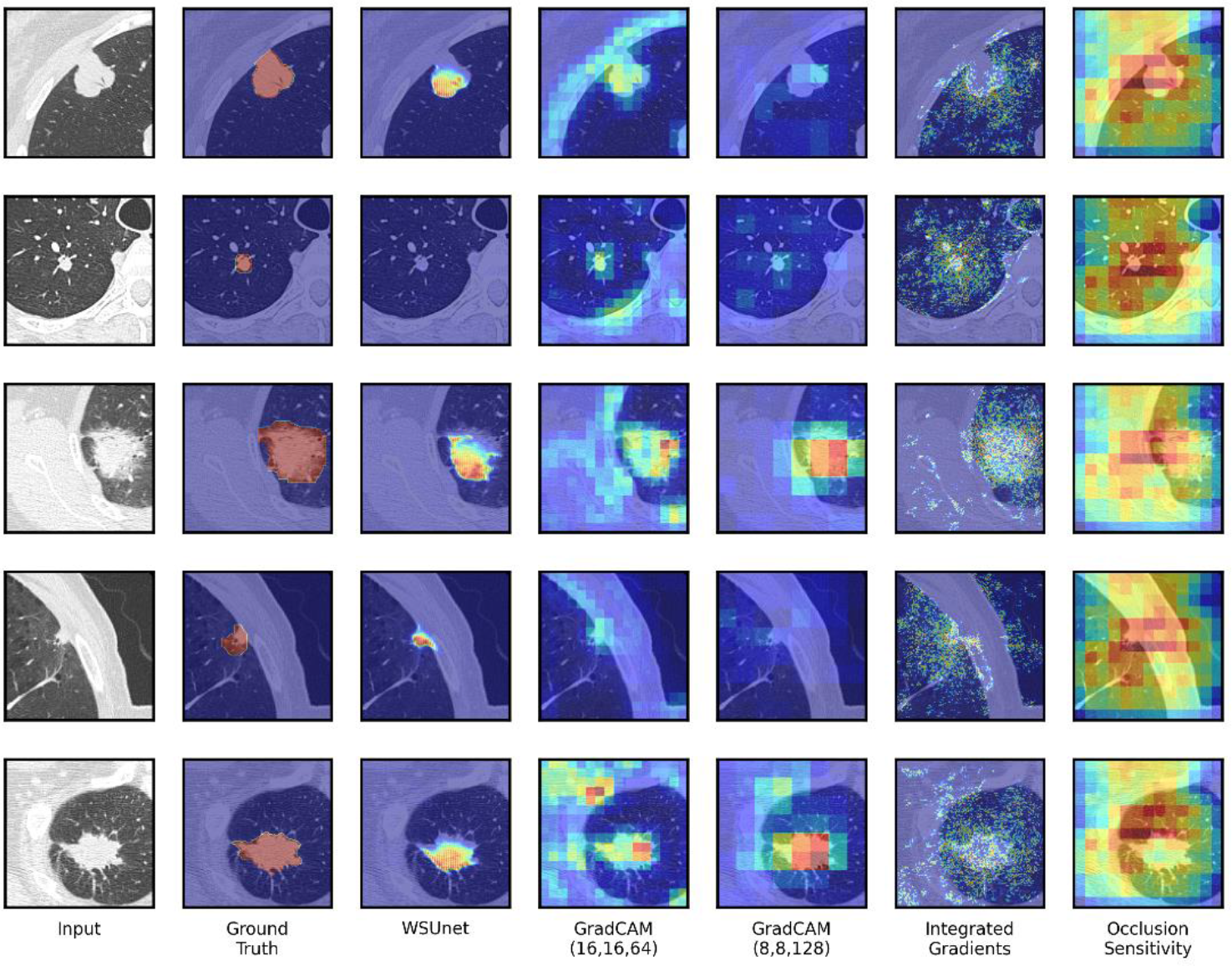
Model explainability heatmaps. The first five positive test images are shown. Models were trained to detect NSCLC tumours at image-level. WSUnet’s heatmap was extracted from the penultimate voxel-level output layer. GradCAM heatmaps were extracted from the seventh (“GradCAM (16,16,64)”) and ninth (“GradCAM (8,8,128)”) convolutional layers. Nearest-neighbour interpolation was applied to map GradCAM, Integrated Gradients and Occlusion Sensitivity heatmap to the input image dimensions. For comparability, methods heatmaps were normalised to the range of minimum and maximum values for the five images.

## DISCUSSION

In this study, we have developed a CNN architecture, WSUnet, which supports voxel-level modelling with image-level labels. By application to NSCLC detection, we have demonstrated that WSUNet may be used to learn voxel-level classification from data labelled only at image-level. Most importantly, we demonstrate that WSUnet explains its image-level predictions at voxel-level in a causally verifiable manner.

WSUnet yielded superior voxel-level discrimination to current model interpretation approaches, both by objective and subjective metrics. WSUnet’s voxel-level output identified the voxels motivating the positive image-level prediction, revealing whether the model attended to the tumour or other confounding features. WSUnet offered a distinct advantage of returning predictions in the domain and range of the voxel-level class probabilities, obviating the need for post-hoc interpolations and transformations. Thus, WSUnet’s voxel-level output could be interpreted directly as a voxel-probability heatmap. Although WSUnet’s voxel-level recall did not challenge the state-of-the art set by fully-supervised NSCLC segmentation models trained under full supervision (Liu et al., 2021), its high precision presents a plausible avenue for object localisation. This result also provides insight into how WSUnet works – if the model detects any part of a tumour, it can deduce that the image is positive. Thus, a positive image-level prediction may be inferred without observing the whole tumour region. Conversely, the whole image must be considered to exclude a tumour. Thus, the model is negatively biased at voxel level. Indeed, it may be observed in **Figure 2** that WSUnet tended to identify the tumour by its posterior aspect.

Although GradCAM predictions localised moderately well to the tumour, their utility was limited by low resolution. Integrated gradients outputs were not locally consistent, such that adjacent voxels typically had dissimilar predictions. Occlusion sensitivity results demonstrated little variance between images. All methods were limited by producing an output which could not be interpreted directly as a voxel-probability map.

It is noted that the ground truth segmentation was performed with access to the entire CT volume, rather than a patch from a single slice. As the inclusion of even a single positive pixel in the patch led to a positive image-level classification, some tumours were difficult to visualise within individual patches. Clinicians annotated several positive test images as “tumour not visible in this image”. In several instances tumours were identifiable only by ground glass changes. Thus, the imperfect image-level classification performance demonstrated by both models may be explained by the difficulty of the test instances. WSUnet provided similar image-level classification performance to SCNN, whilst delivering significant improvements in terms of interpretability.

WSUnet is a CNN which returns both an image-level decision and a voxel-level saliency map which caused the decision. This development facilitates model inspection, debugging, reliability testing, inference and pathobiological discovery. The approach differs significantly from current model inspection methods, as the image-level prediction is simply the maximal voxel-level probability. Consequently, voxel-level outputs are easily interpretable as class probabilities and provide a causally verifiable explanation for the image-level decision. Although the WSUnet architecture implemented here generates a scalar output representing binary classifications, multi-class configurations are theoretically achievable with corresponding architectures. It is also conceivable that higher-order cognitive functions encoded in internal convolutional layers may be inspected with architectural variations of the WSUnet.

This retrospective study included model evaluation on multi-centre data which was geographically distinct from training data. Methods were compared under both objective and subjective metrics. The WSUnet model successfully learned voxel-level classification functions from a dataset of image-level labelled data. WSUnet’s voxel-level output is demonstrated to be causally linked to the image-level output. WSUnet’s output has the domain and distribution of the class labels, and it may be interpreted directly as a set of voxel-level class probability estimates. The class distribution in this proof-of-concept study was approximately balanced at image-level and moderately imbalanced at voxel level – convergence of weak-learners may be less reliable in severely imbalanced data. The implementations in this proof-of-concept study were rudimentary - superior performance should be expected with optimised training routines and model hyperparameters.

## CONCLUSIONS

In conclusion, this study demonstrated that a simple extension of the traditional FCN yielded an image-level classifier whose penultimate layer provided a causal explanation of its decision at voxel level. This method jointly addresses the related problems of weakly supervised semantic segmentation and convolutional neural network explainability. Thus, WSUnet may be used to improve the clinical reliability of CNN image classifiers. An exciting prospect for future research is the application of the weakly supervised FCN paradigm to other biomedical domains. For example, in disease modelling from electronic health records, it may be possible to identify the terms, phrases or paragraphs which support the predicted diagnosis.

## Supporting information

The clinician preference survey is provided in Supplementary Data 1

## Abbreviations

18F-FDG PET: 18-fluorine fluorodeoxyglucose positron emission tomography
AUC: area under receiver operator characteristic curve
AUPR: area under precision recall curve
CNN: convolutional neural network
CT: computed tomography
FCN: fully convolutional neural network
NSCLC: non-small cell lung cancer

## Acknowledgements

The authors would like to thank Aerts et al., Bakr et al. and The Cancer Imaging Archive for providing the study data.

## DATA AVAILABILITY STATEMENT

This study involved no data collected by study authors. All datasets used in this study are publicly available from The Cancer Imaging Archive (Aerts et al., 2019, 2014; Bakr et al., 2017; Clark et al., 2013). All code required to reproduce the findings of this study is provided at github.com/robertoshea/wsss.

## AUTHOR’S CONTRIBUTIONS

Conceptualisation: Robert O’Shea.

Methodology: Robert O’Shea, Carolyn Horst, Thubeena Manickavasagar.

Data curation: Robert O’Shea. Formal analysis: Robert O’Shea.

Software: Robert O’Shea.

Validation: Carolyn Horst, Thubeena Manickavasagar, Daniel Hughes, James Cusack, Vicky Goh.

Funding acquisition: Vicky Goh.

Supervision: Sophia Tsoka, Gary Cook, Vicky Goh.

Writing (original draft): Robert O’Shea.

Writing (review & editing): Robert O’Shea, Carolyn Horst, Thubeena Manickavasagar, Daniel Hughes, James Cusack, Gary Cook, Sophia Tsoka, Vicky Goh.

