## Supplementary material for "Weakly supervised Unet: an image classifier which learns to explain itself": The clinician preference survey is provided in Supplementary Data 1

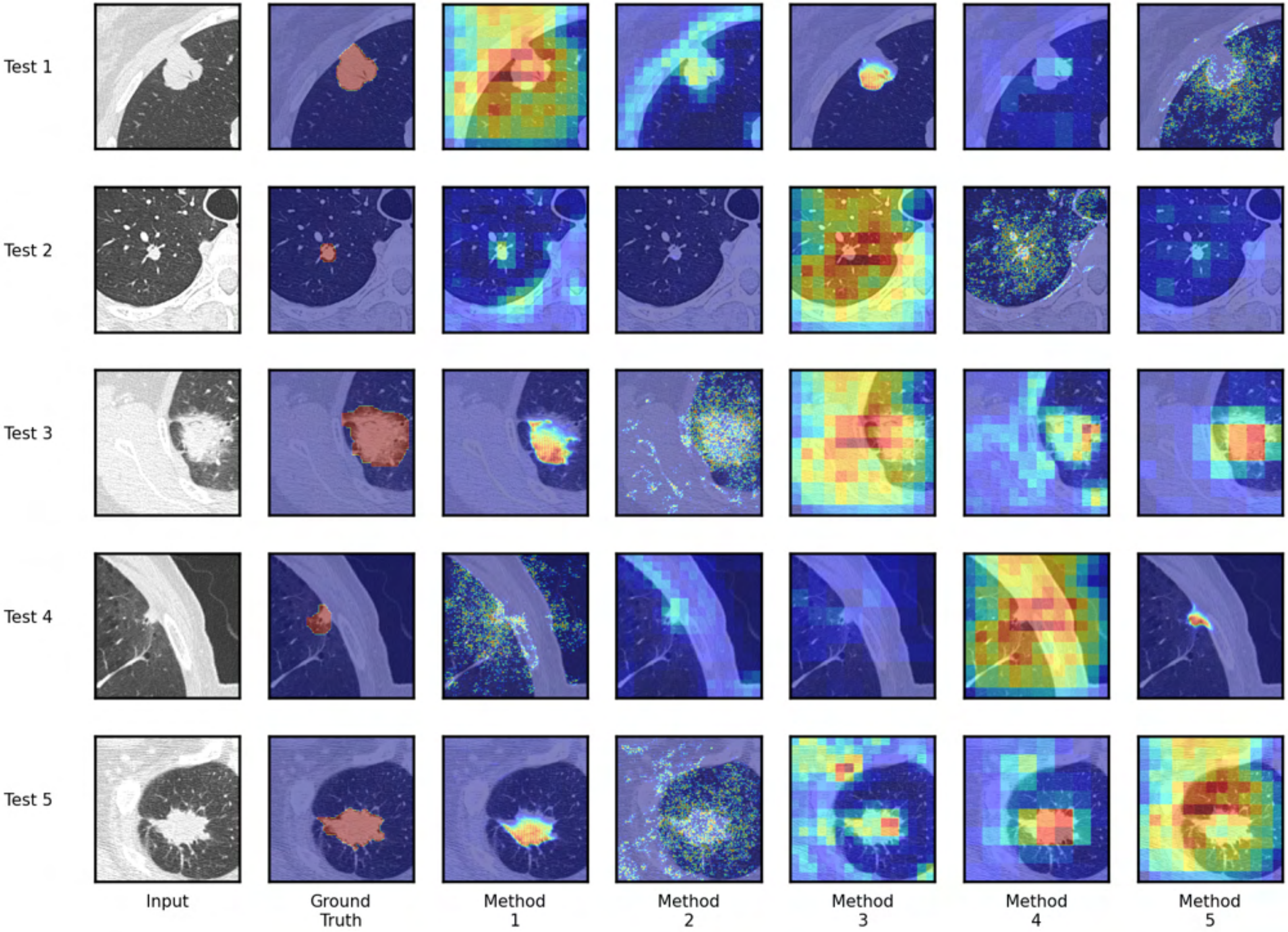
Test 1
Test 2
Test 3
Test 4
Test 5
Input
Ground Truth
Method 1
Method 2
Method 3
Method 4
Method 5

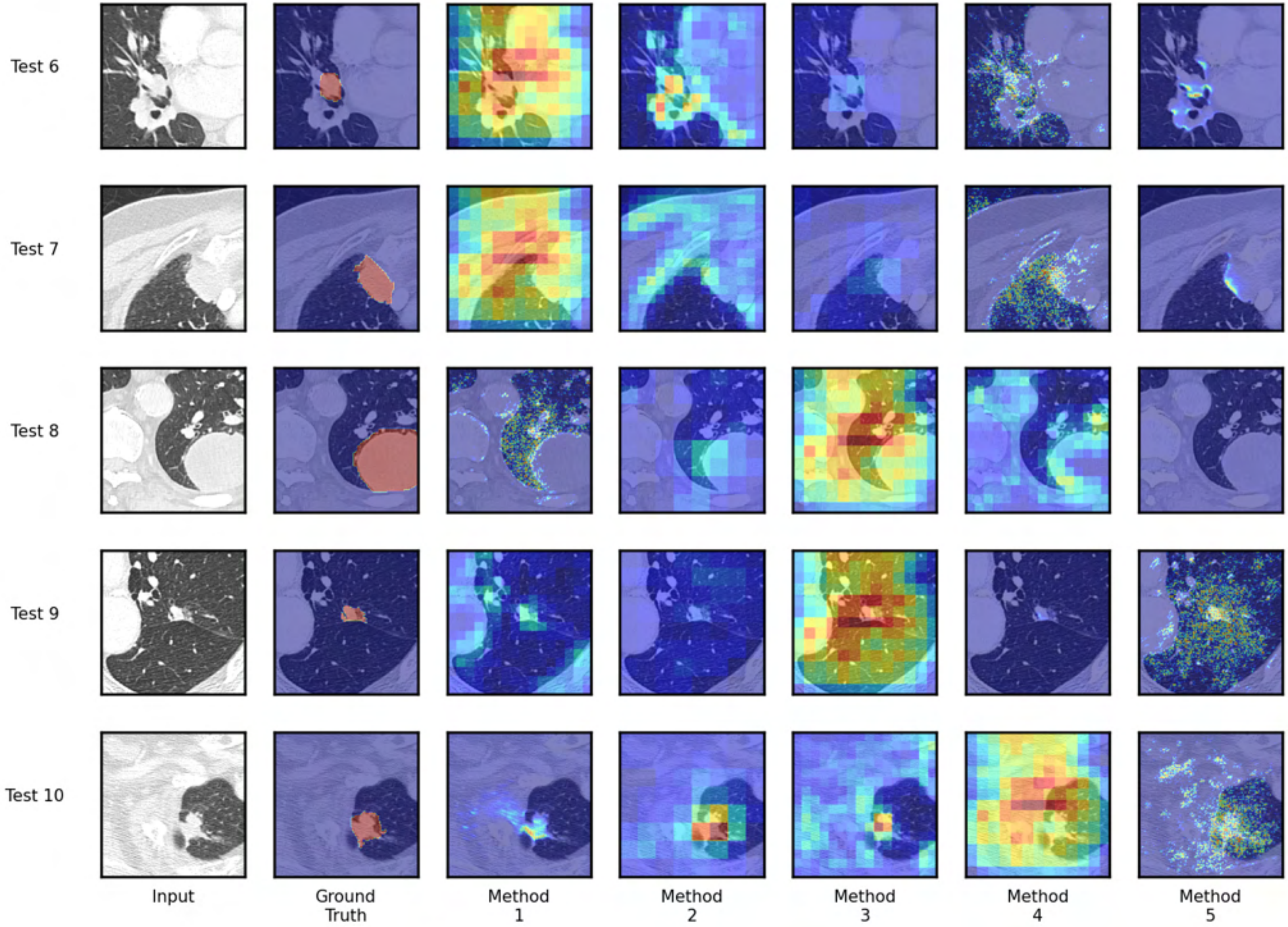

Test 6
Test 7
Test 8
Test 9
Test 10
Input
Ground Truth
Method 1
Method 2
Method 3
Method 4
Method 5

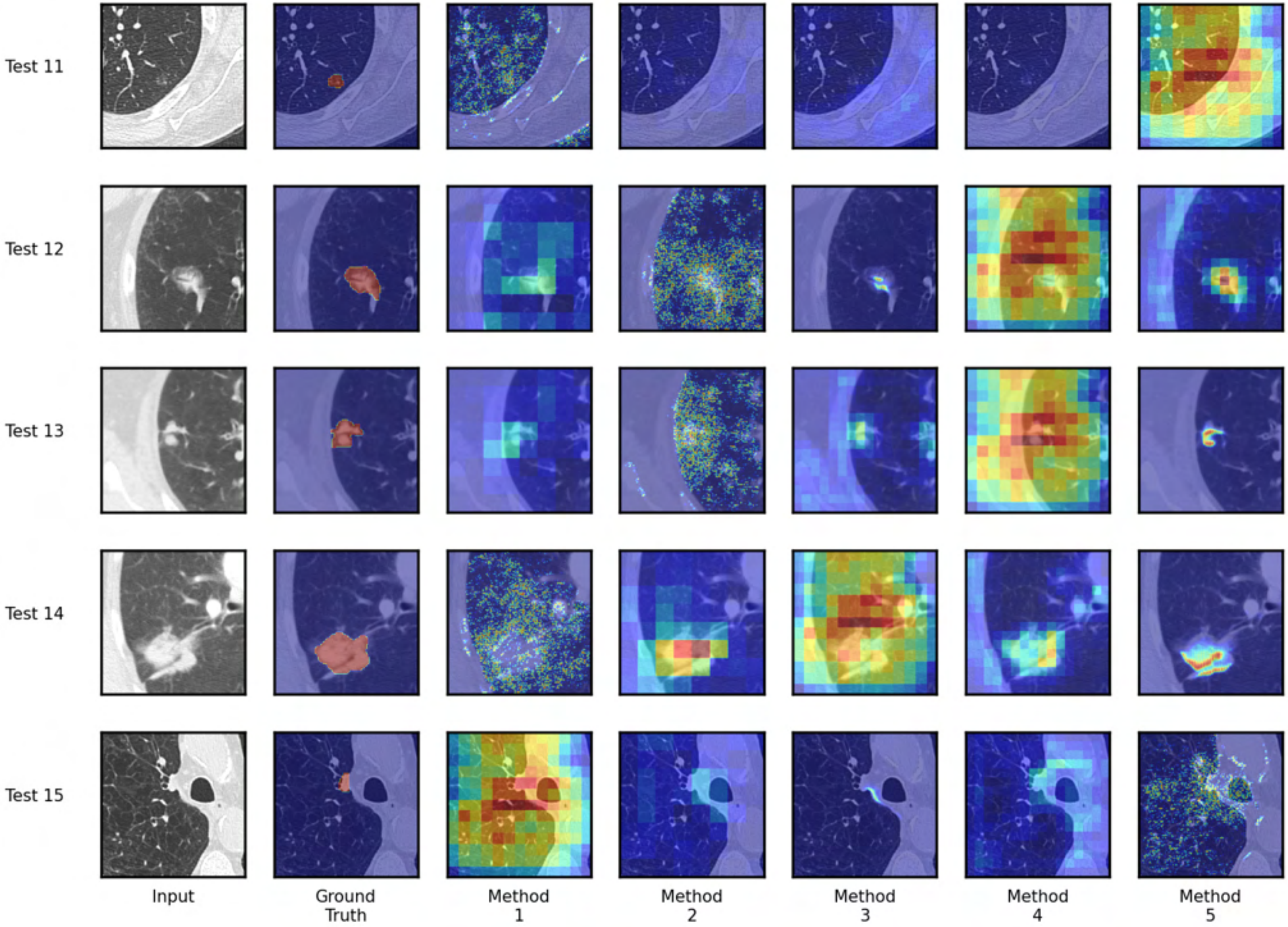
Test 11
Test 12
Test 13
Test 14
Test 15
Input
Ground Truth
Method 1
Method 2
Method 3
Method 4
Method 5

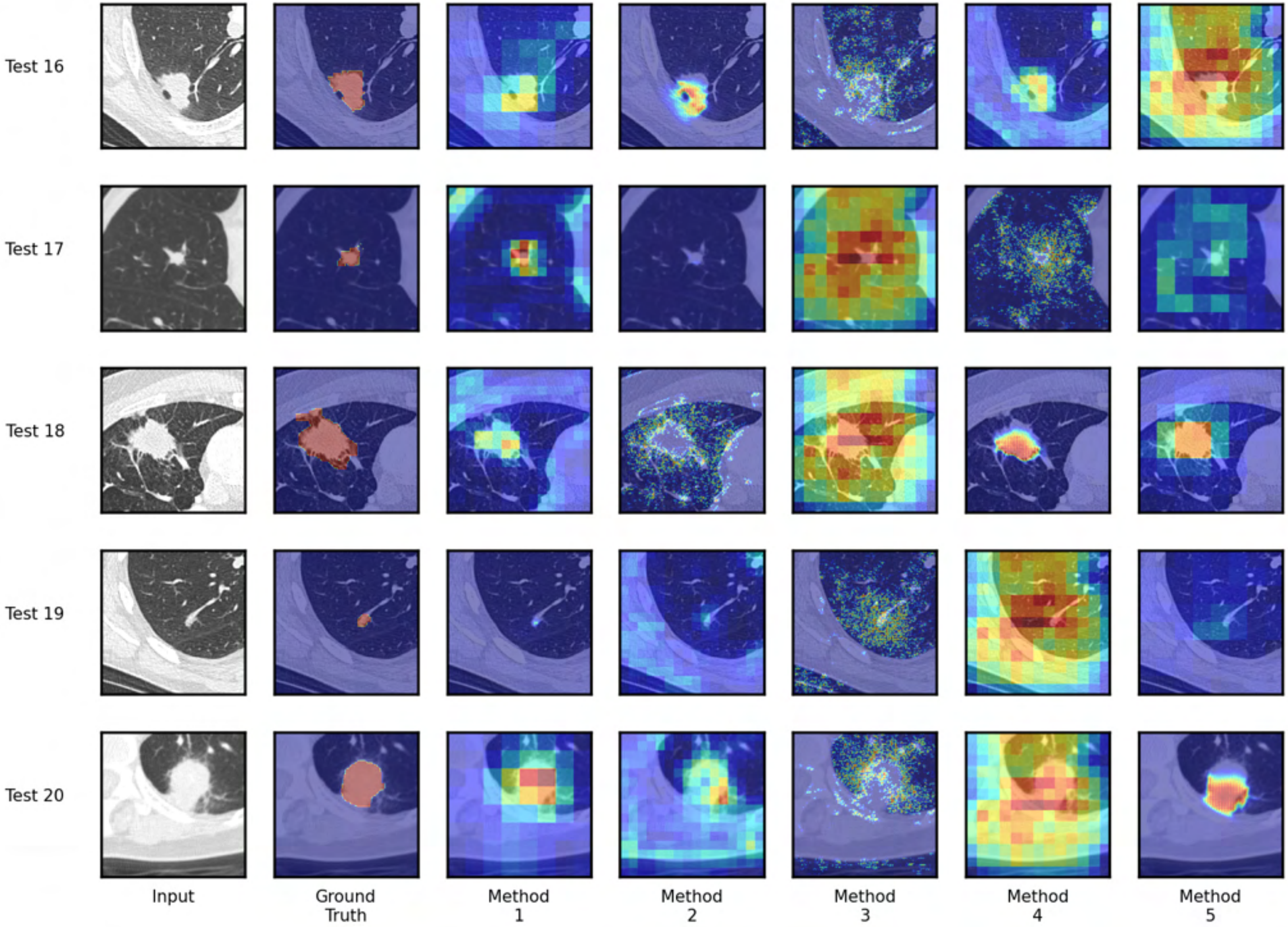
Test 16
Test 17
Test 18
Test 19
Test 20
Input
Ground Truth
Method 1
Method 2
Method 3
Method 4
Method 5

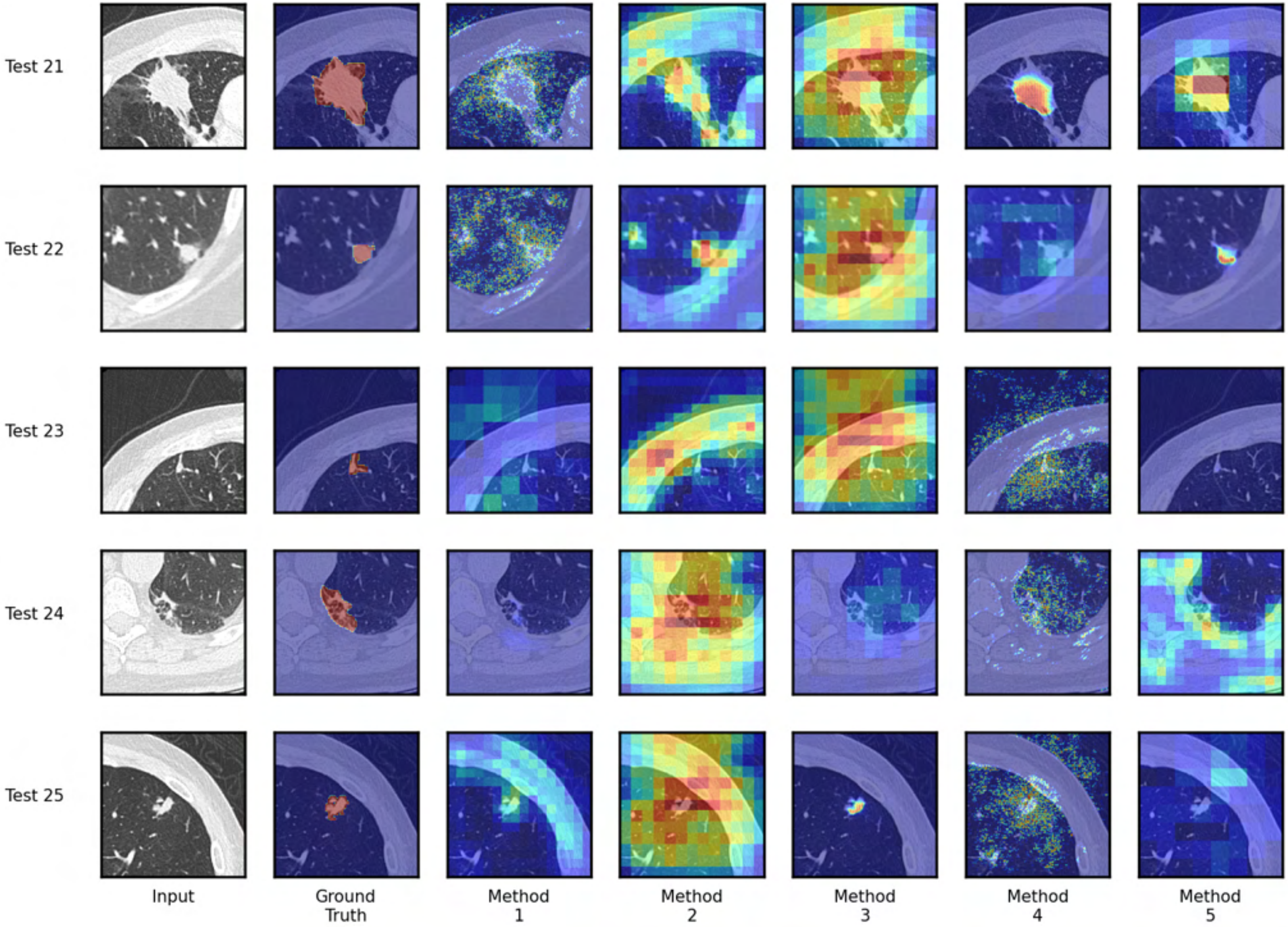
Test 21
Test 22
Test 23
Test 24
Test 25
Input
Ground Truth
Method 1
Method 2
Method 3
Method 4
Method 5

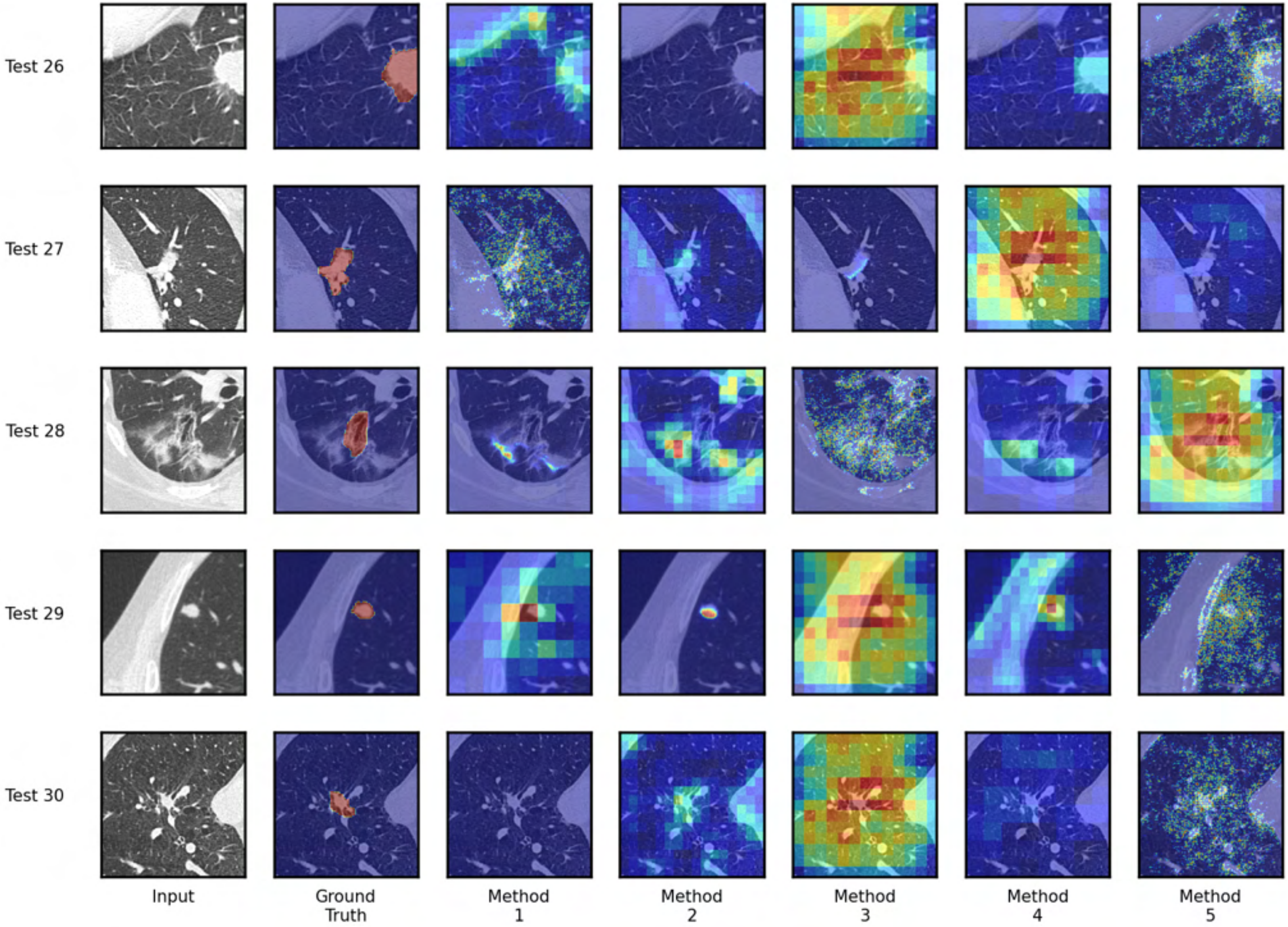

Test 26
Test 27
Test 28
Test 29
Test 30
Input
Ground Truth
Method 1
Method 2
Method 3
Method 4
Method 5

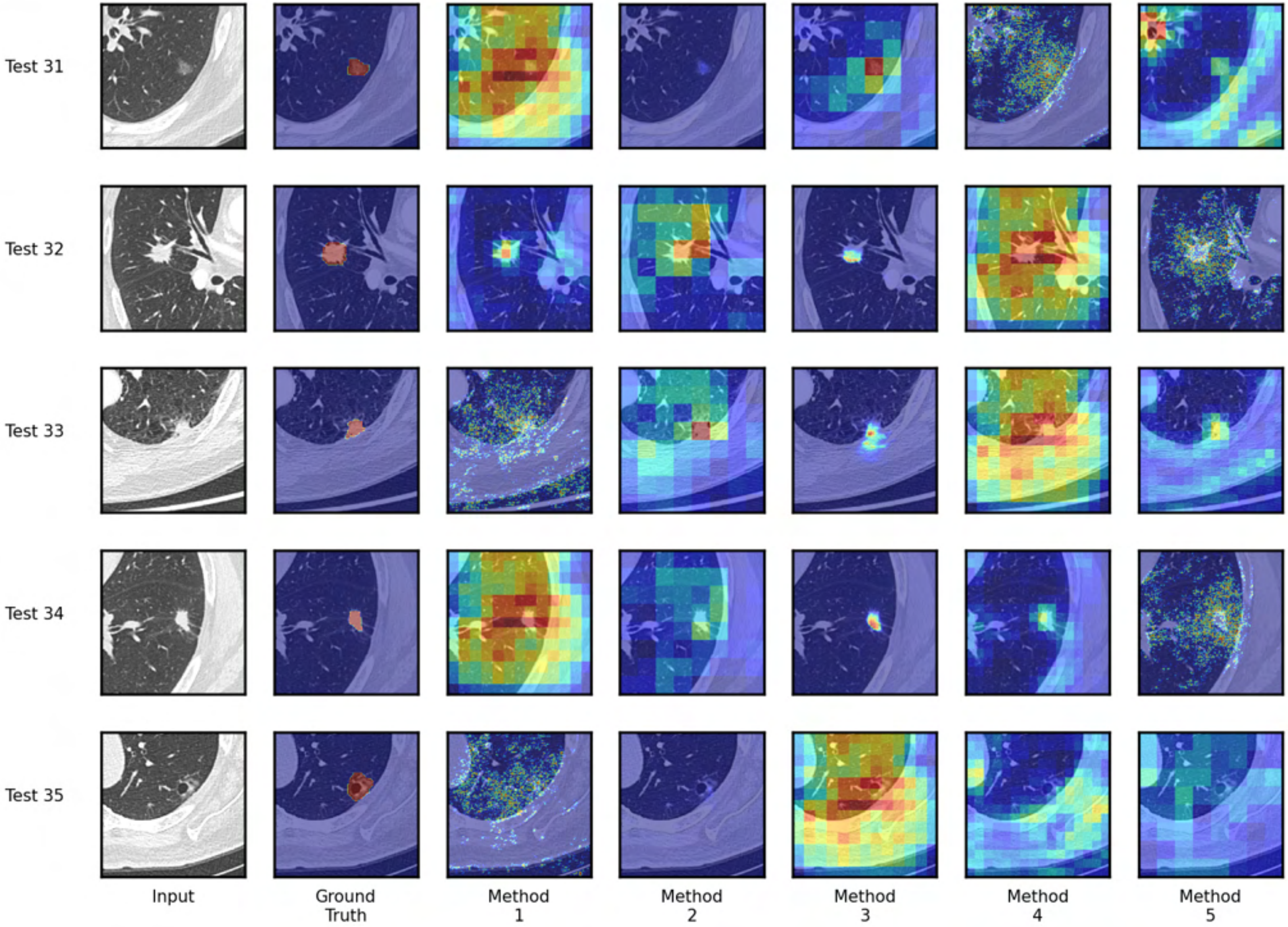

Test 31
Test 32
Test 33
Test 34
Test 35
Input
Ground Truth
Method 1
Method 2
Method 3
Method 4
Method 5

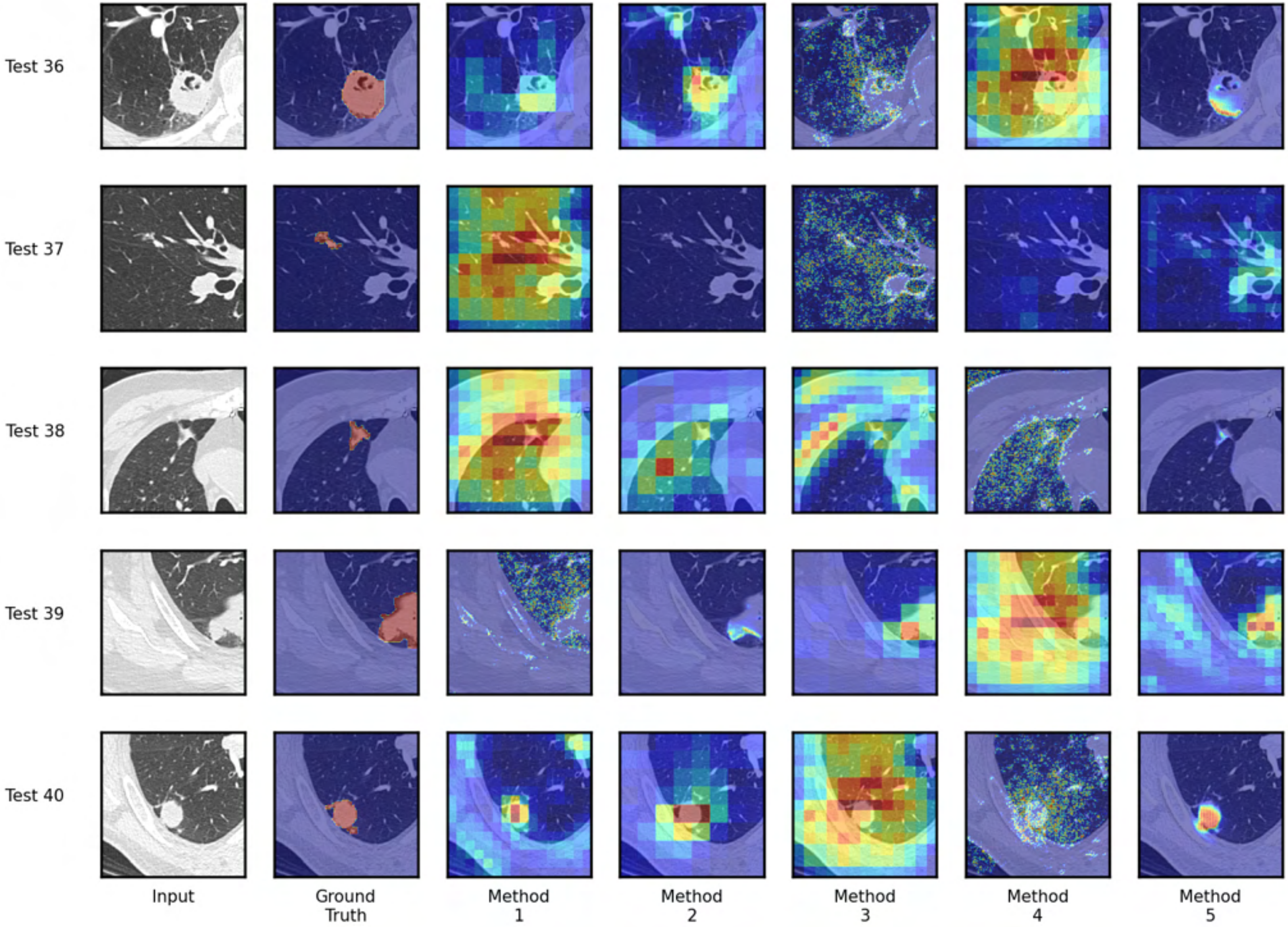
Test 36
Test 37
Test 38
Test 39
Test 40
Input
Ground Truth
Method 1
Method 2
Method 3
Method 4
Method 5

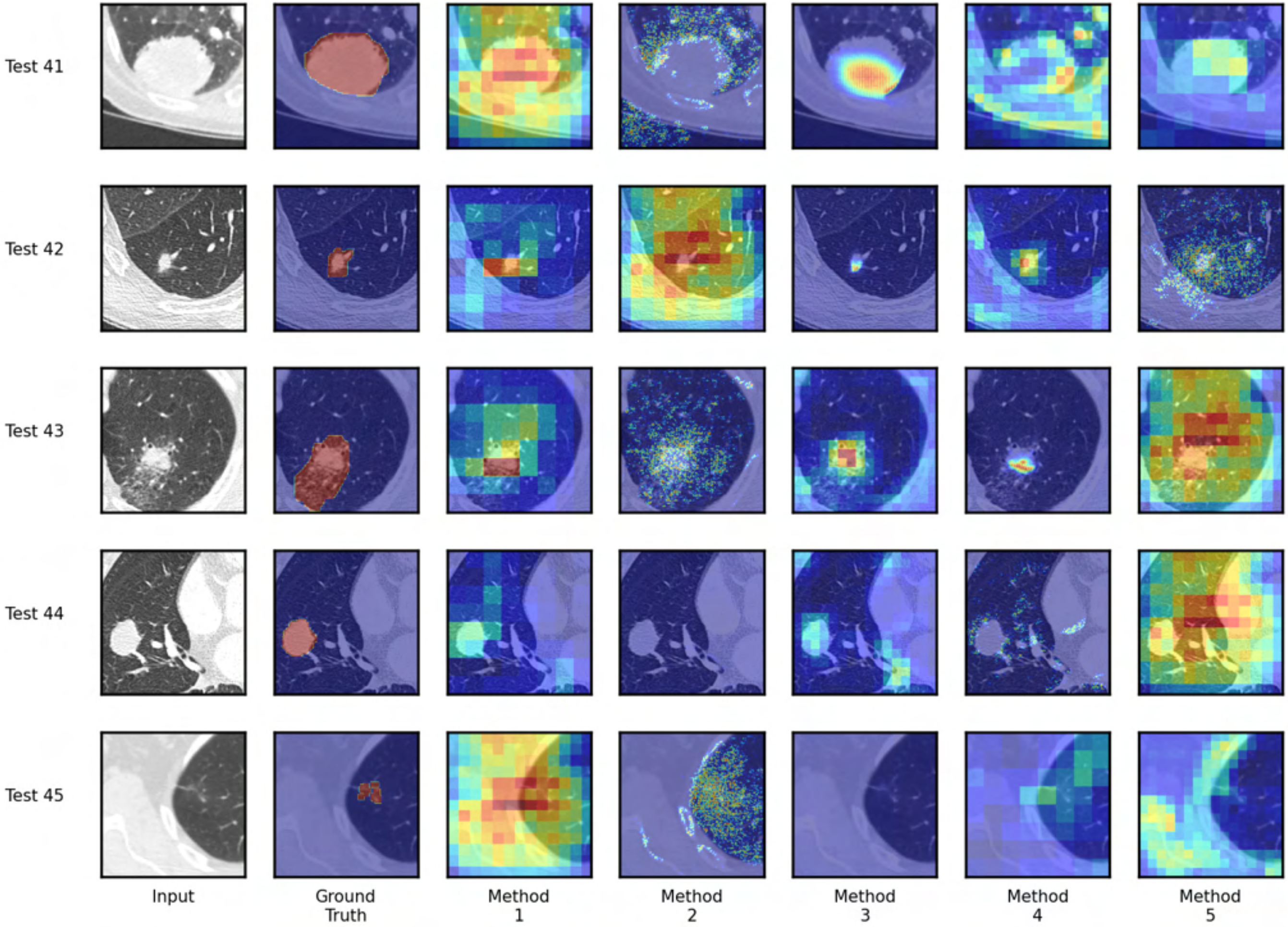
Test 41
Test 42
Test 43
Test 44
Test 45
Input
Ground Truth
Method 1
Method 2
Method 3
Method 4
Method 5

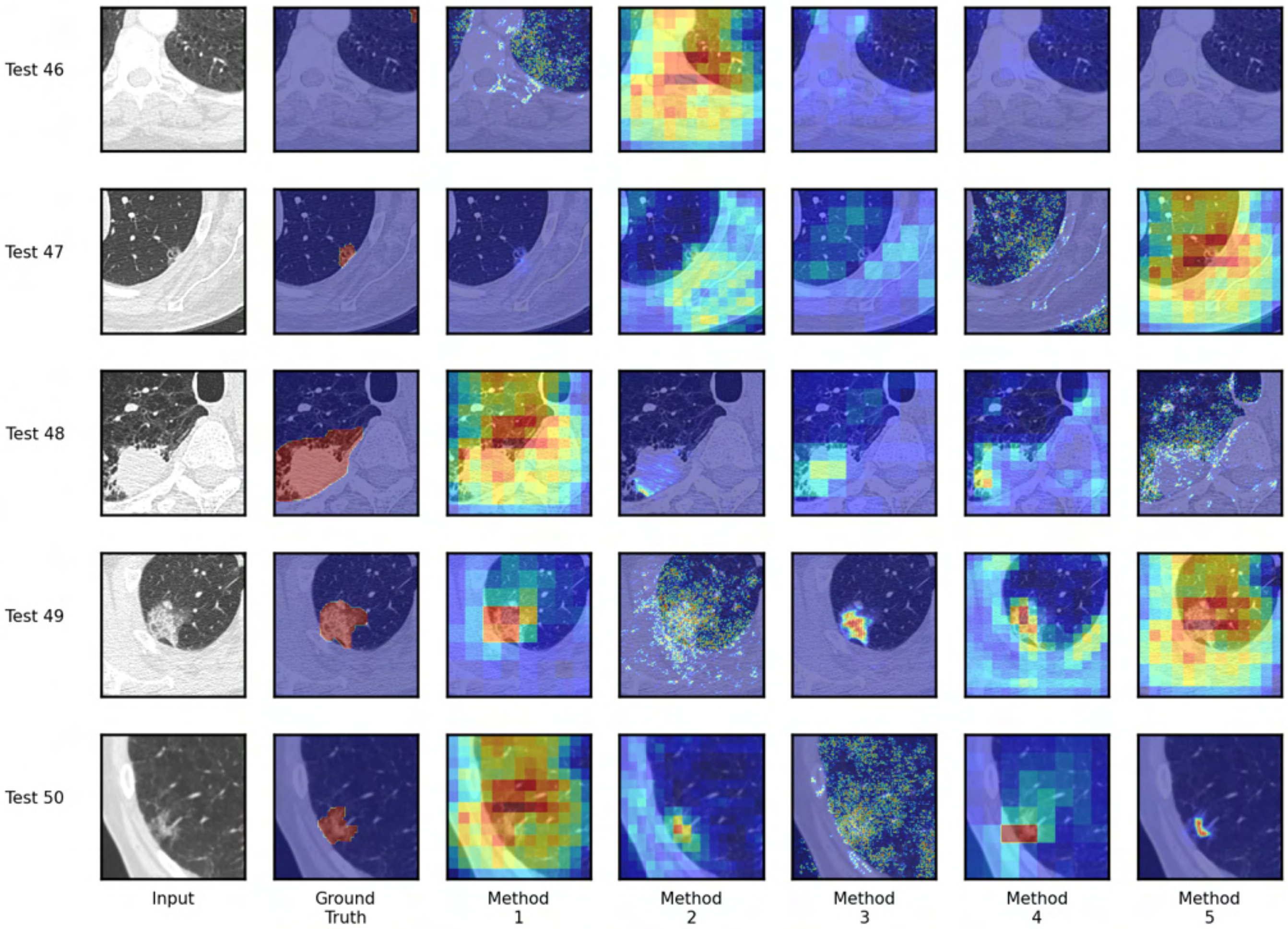
Test 46
Test 47
Test 48
Test 49
Test 50
Input
Ground Truth
Method 1
Method 2
Method 3
Method 4
Method 5

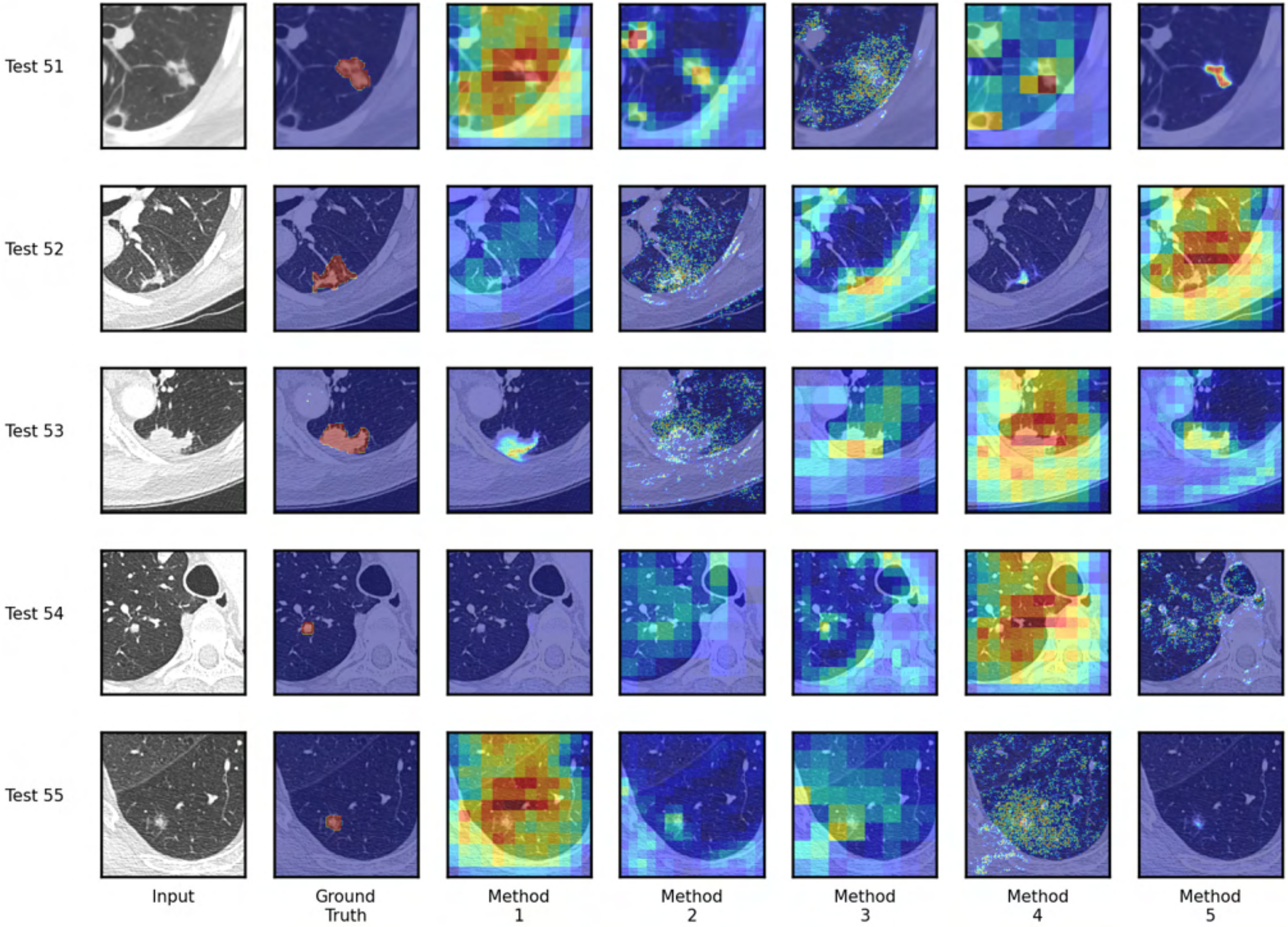
Test 51
Test 52
Test 53
Test 54
Test 55
Input
Ground Truth
Method 1
Method 2
Method 3
Method 4
Method 5

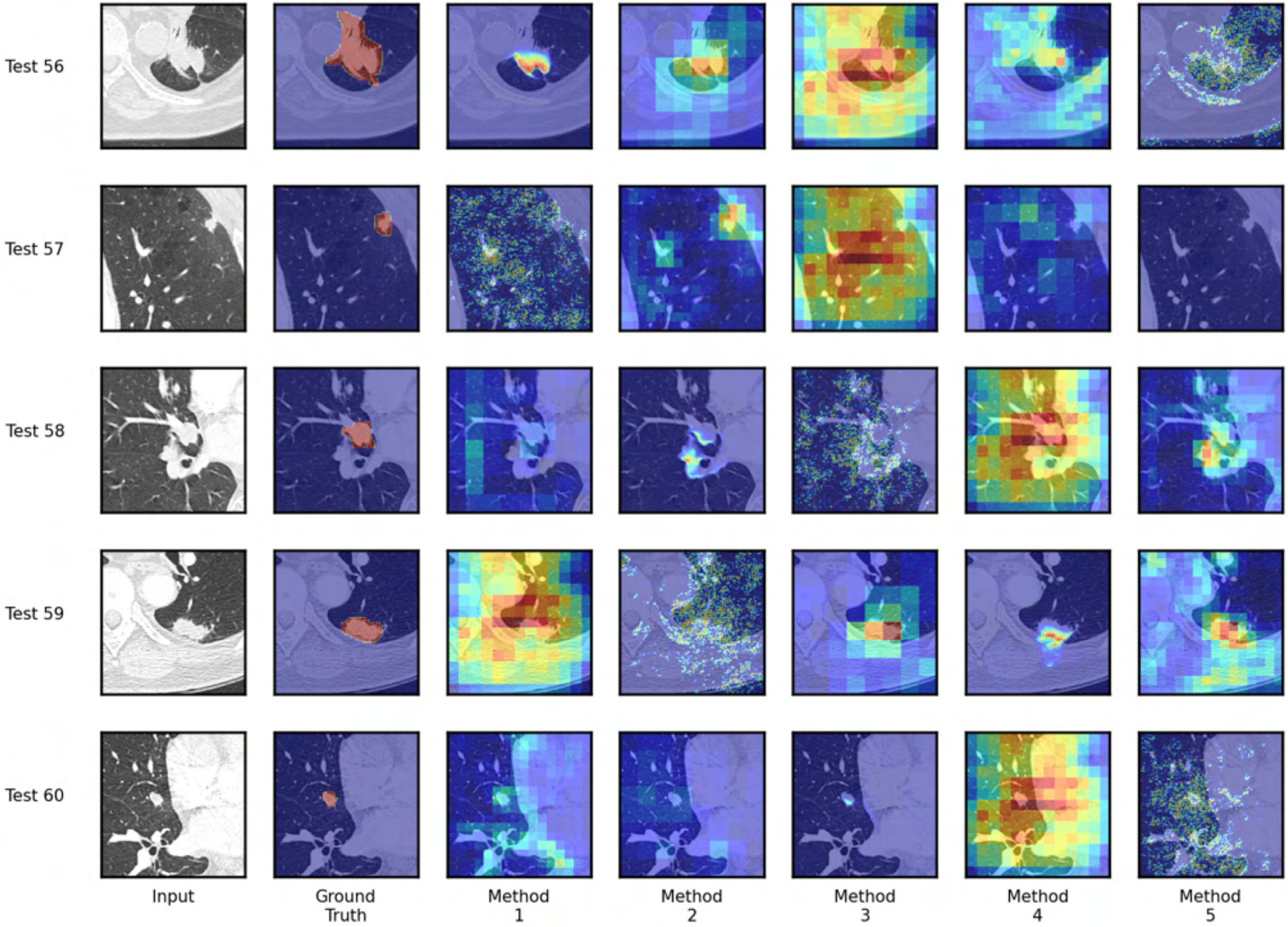

Test 56
Test 57
Test 58
Test 59
Test 60
Input
Ground Truth
Method 1
Method 2
Method 3
Method 4
Method 5

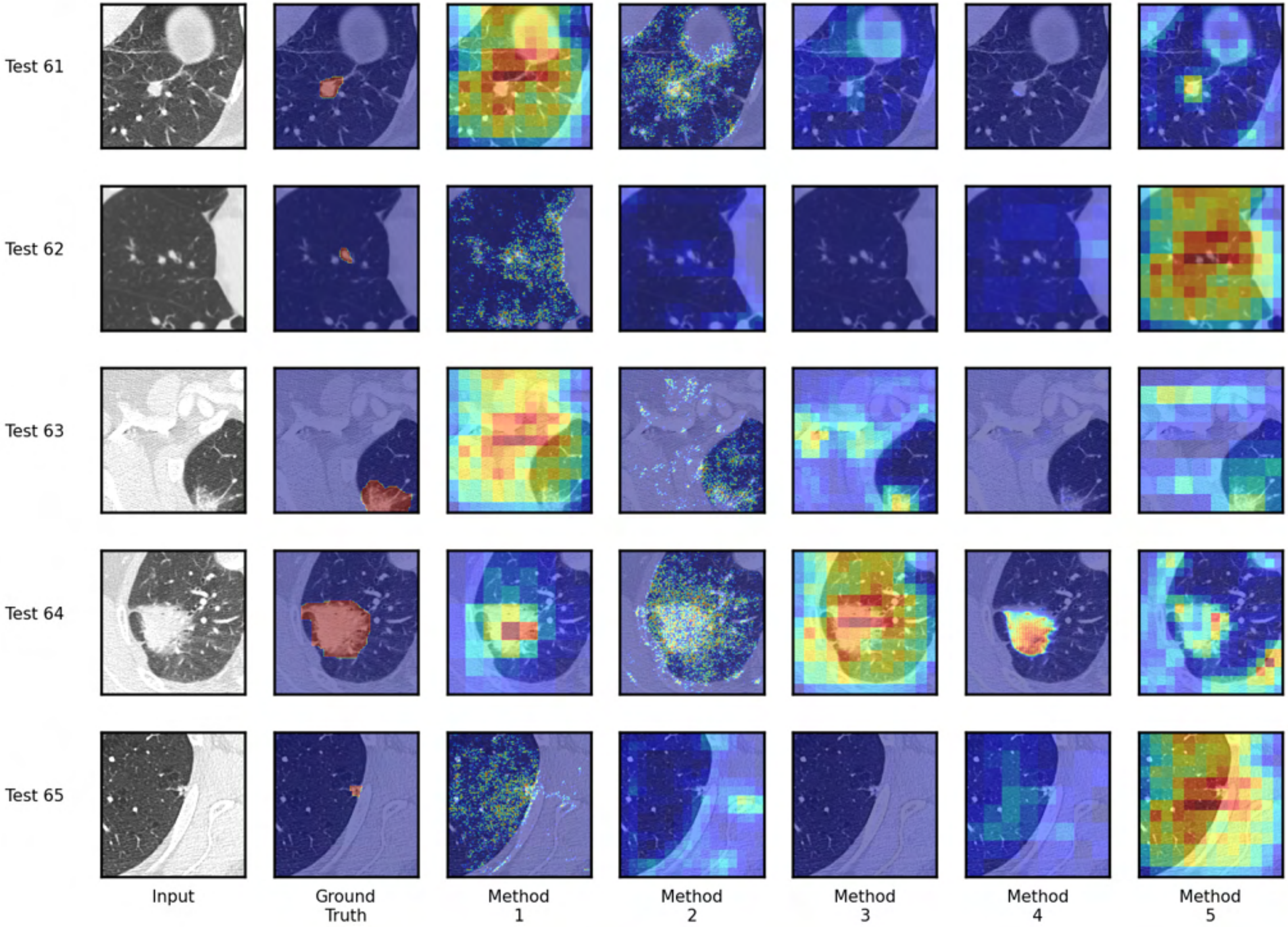
Test 61
Test 62
Test 63
Test 64
Test 65
Input
Ground Truth
Method 1
Method 2
Method 3
Method 4
Method 5

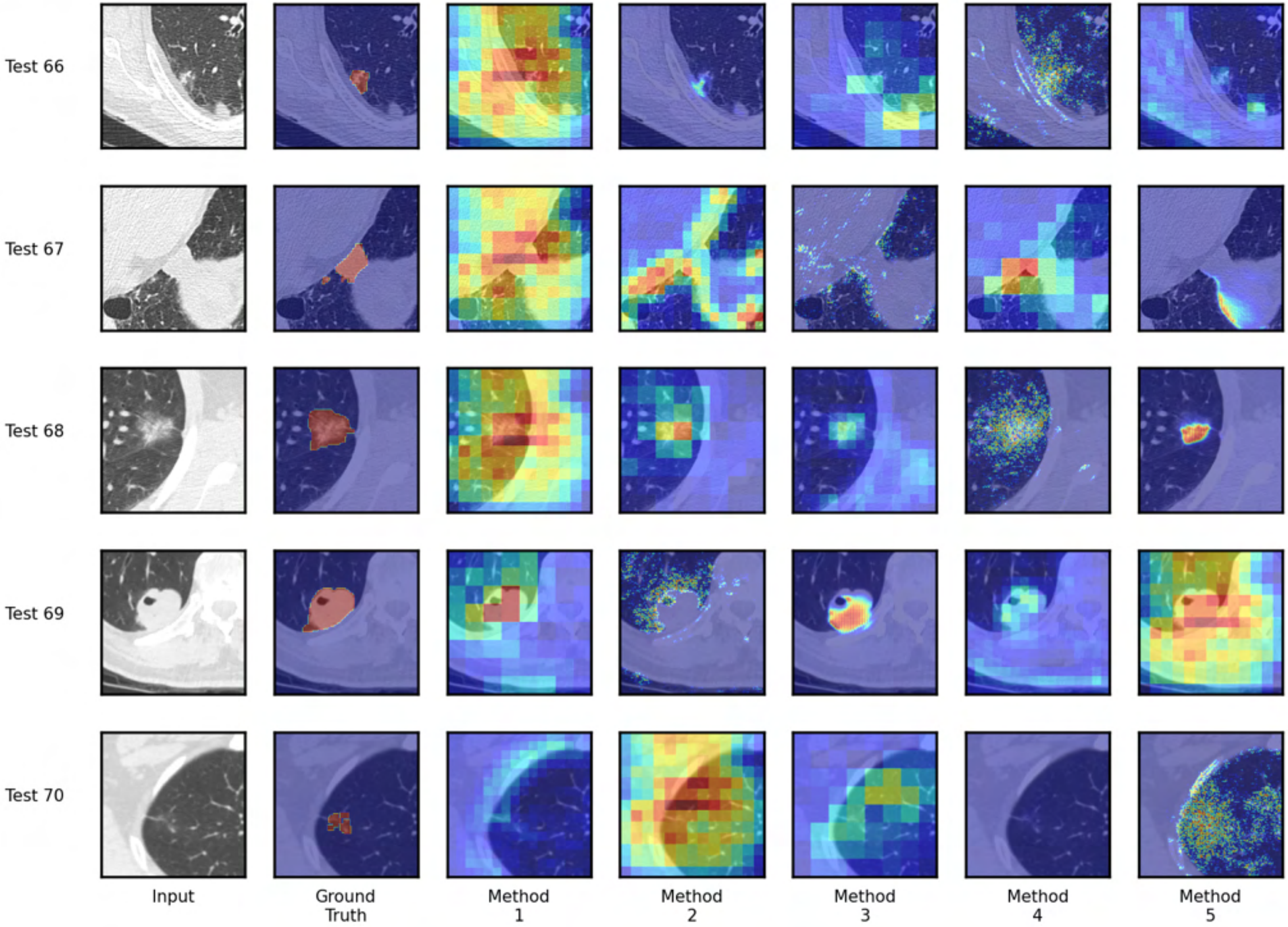
Test 66
Test 67
Test 68
Test 69
Test 70
Input
Ground Truth
Method 1
Method 2
Method 3
Method 4
Method 5

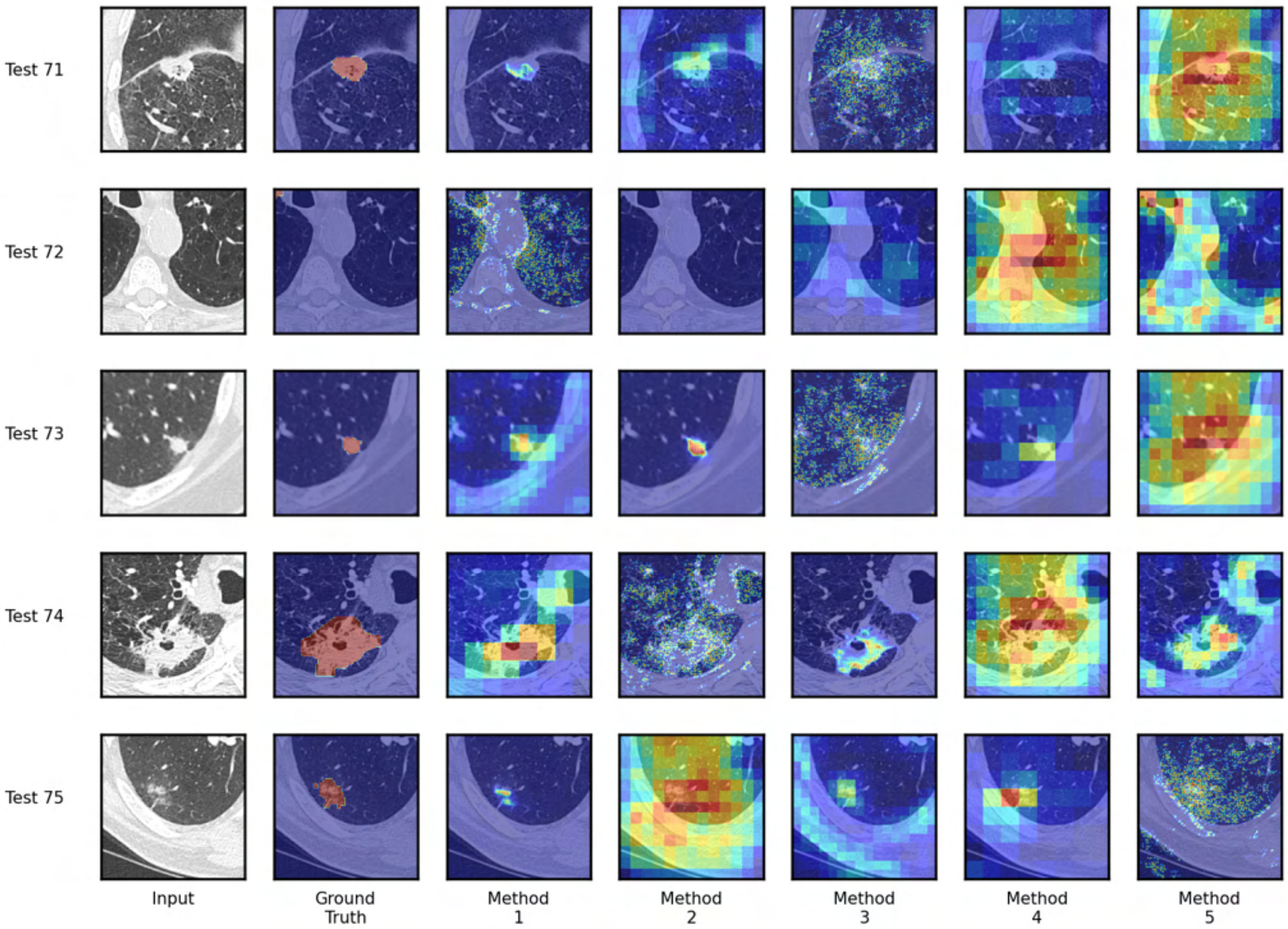
Test 71
Test 72
Test 73
Test 74
Test 75
Input
Ground Truth
Method 1
Method 2
Method 3
Method 4
Method 5

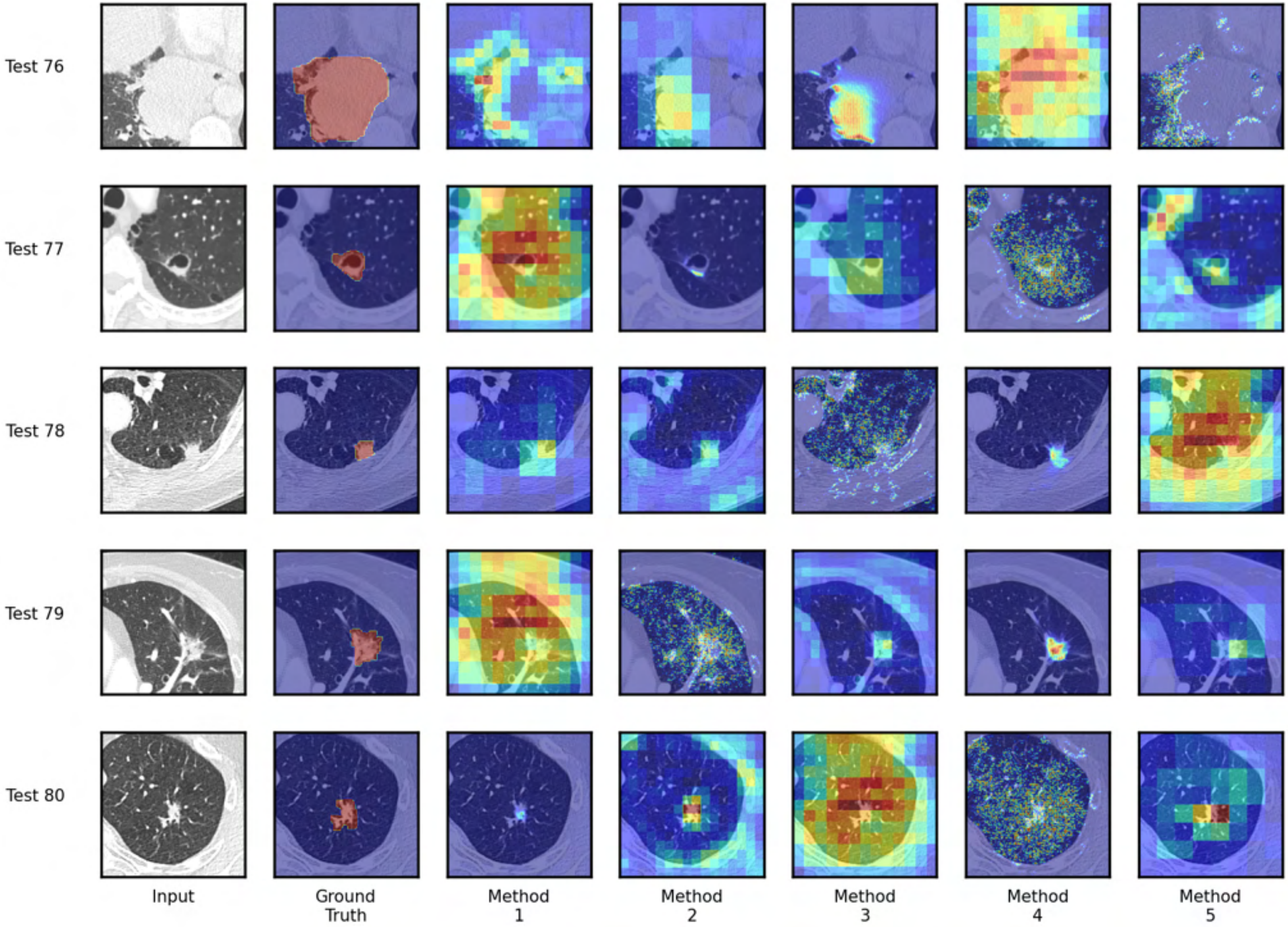
Test 76
Test 77
Test 78
Test 79
Test 80
Input
Ground Truth
Method 1
Method 2
Method 3
Method 4
Method 5

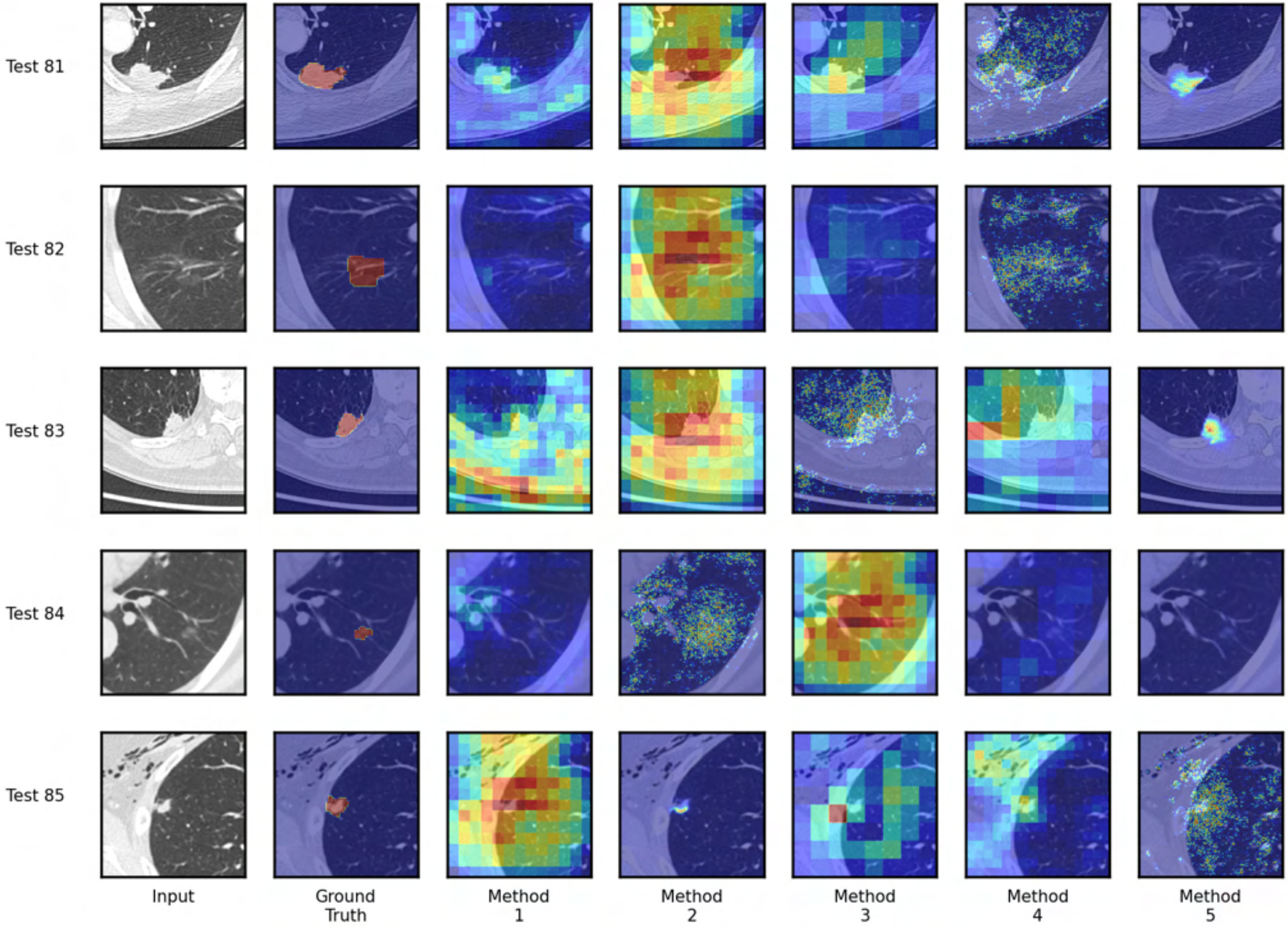

Test 81
Test 82
Test 83
Test 84
Test 85
Input
Ground Truth
Method 1
Method 2
Method 3
Method 4
Method 5

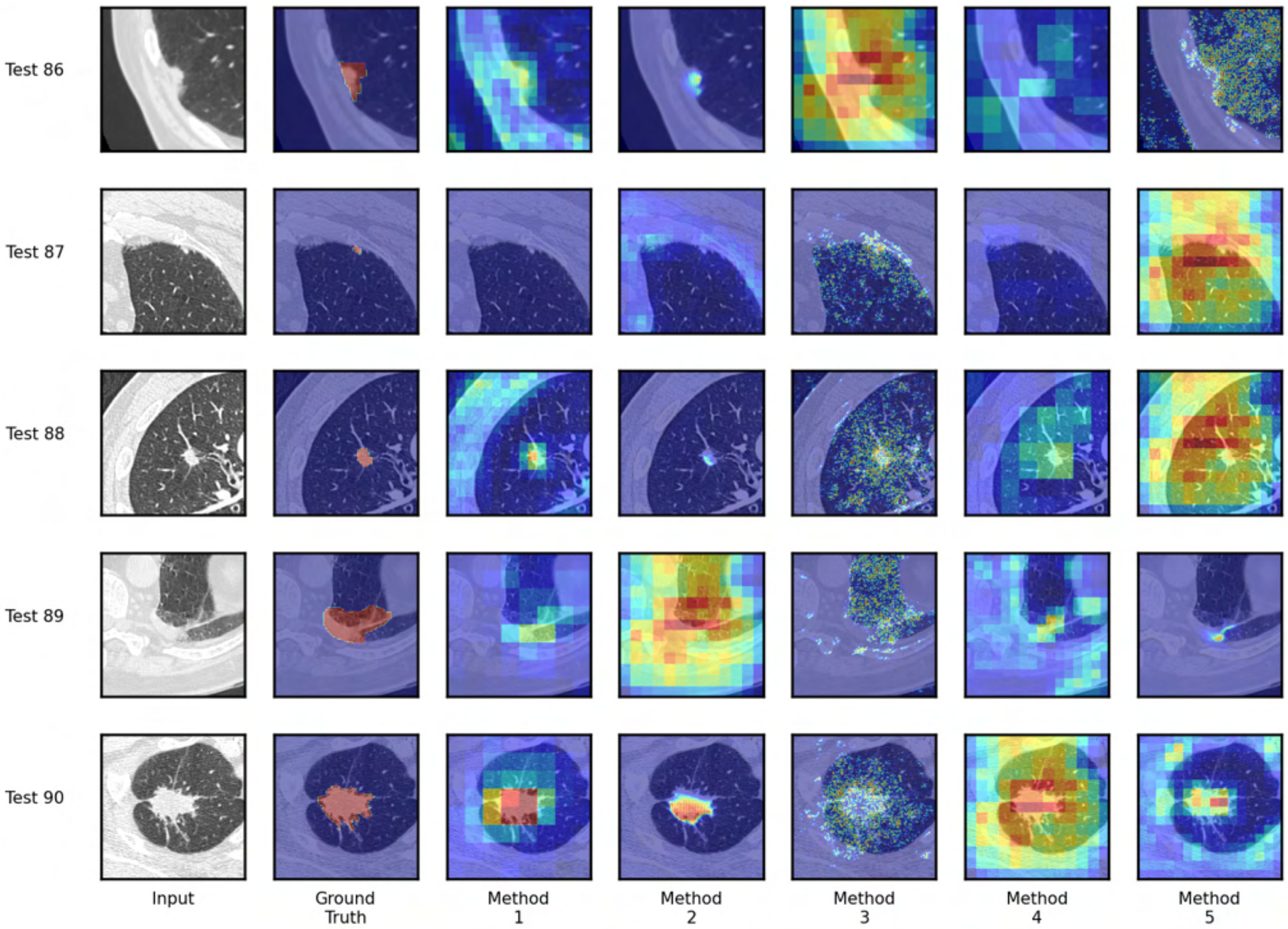

Test 86
Test 87
Test 88
Test 89
Test 90
Input
Ground Truth
Method 1
Method 2
Method 3
Method 4
Method 5

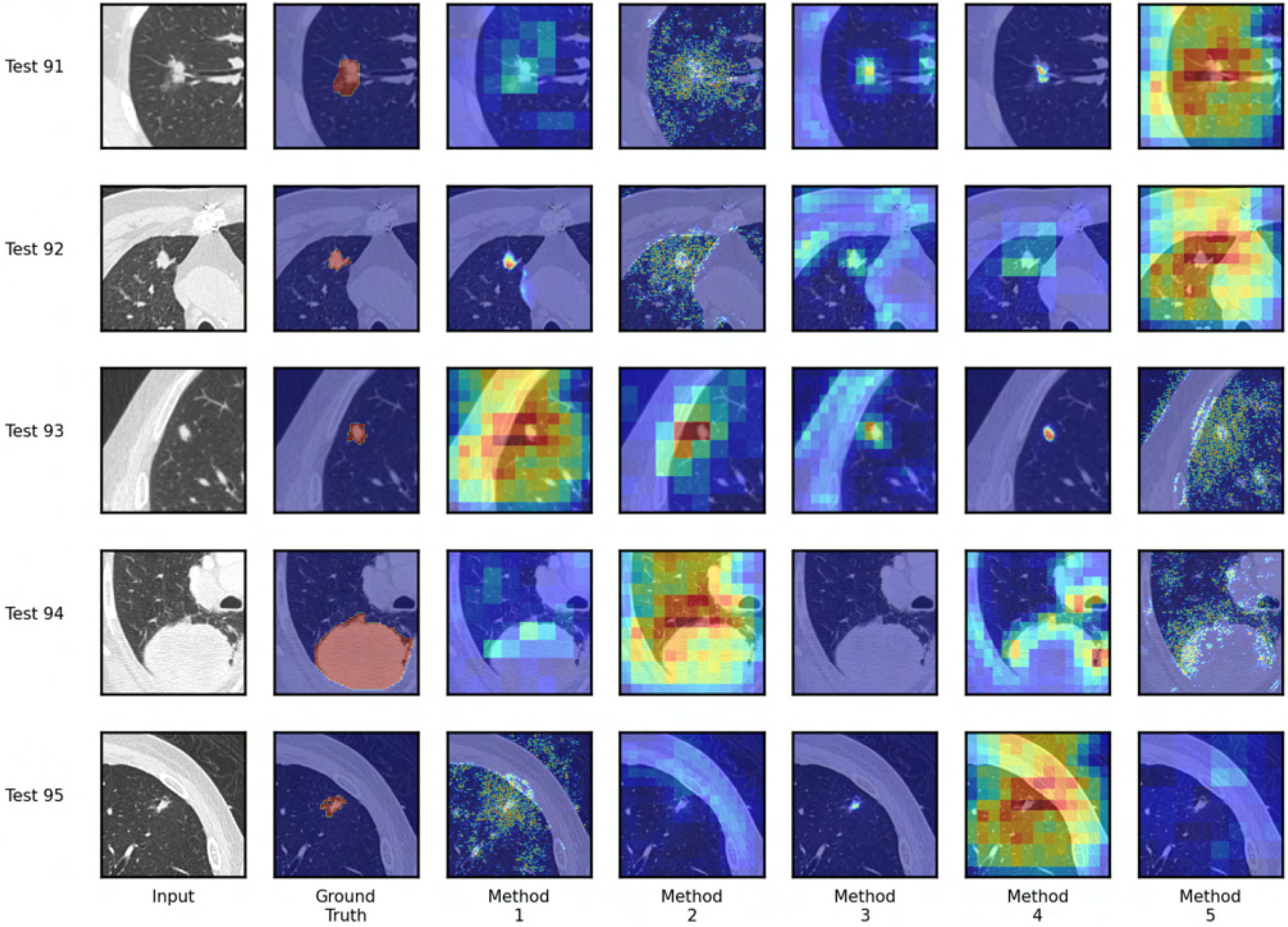
Test 91
Test 92
Test 93
Test 94
Test 95
Input
Ground Truth
Method 1
Method 2
Method 3
Method 4
Method 5

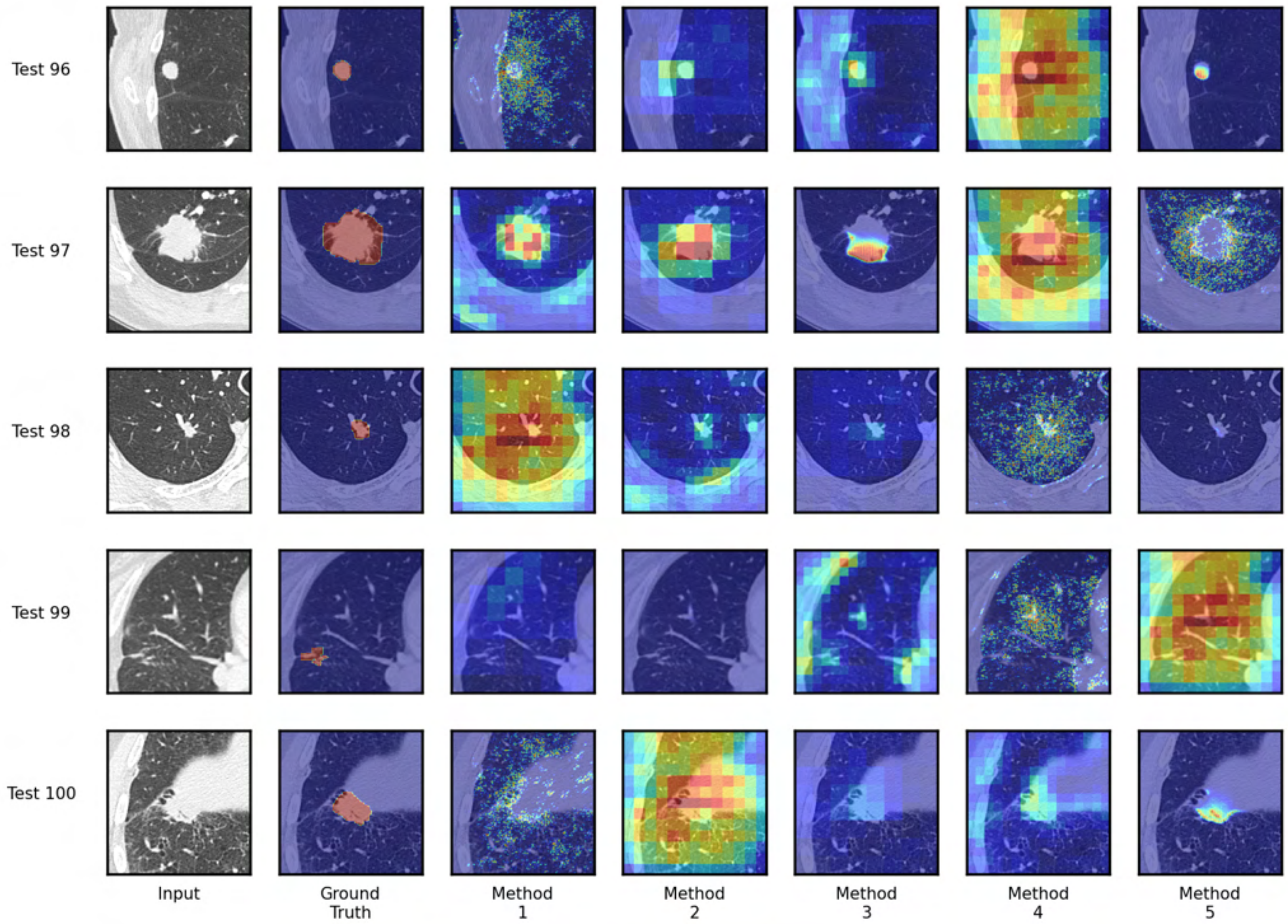
Test 96
Test 97
Test 98
Test 99
Test 100
Input
Ground Truth
Method 1
Method 2
Method 3
Method 4
Method 5
